# Stimulation parameters shape effective connectivity pathways: insights from microstate analysis on TMS-evoked potentials

**DOI:** 10.1101/2024.12.18.629242

**Authors:** Delia Lucarelli, Giacomo Guidali, Dominika Sulcova, Agnese Zazio, Natale Salvatore Bonfiglio, Antonietta Stango, Guido Barchiesi, Marta Bortoletto

## Abstract

Transcranial magnetic stimulation (TMS)-evoked potentials (TEPs) represent an innovative measure for examining brain connectivity and developing biomarkers of psychiatric conditions. Minimising TEP variability across studies and participants, which may stem from methodological choices, is therefore vital. By combining classic peak analysis and microstate investigation, we tested how TMS pulse waveform and current direction may affect effective connectivity when targeting the primary motor cortex (M1). We aim to disentangle whether changing these parameters affects the degree of activation of the same neural circuitry or may lead to changes in the pathways through which the induced activation spreads.

Thirty-two healthy participants underwent a TMS-EEG experiment in which the pulse waveform (monophasic, biphasic) and current direction (posterior-anterior, anterior-posterior, latero-medial) were manipulated. We assessed the latency and amplitude of M1-TEP components and employed microstate analyses to test differences in topographies.

Results revealed that TMS parameters strongly influenced M1-TEP components’ amplitude but had a weaker role over their latencies. Importantly, microstate analysis showed that the current direction in monophasic stimulations changed the pattern of evoked microstates at the early TEP latencies, as well as their duration and the overall amount of activated brain resources associated.

This study shows that the current direction of monophasic pulses may modulate cortical sources contributing to TEP signals, activating neural populations and cortico-cortical paths more selectively. Biphasic stimulation reduces the variability associated with current direction and may be better suited when TMS targeting is blind to anatomical information.

## 1. INTRODUCTION

The combination of transcranial magnetic stimulation and electroencephalography (TMS-EEG) allows the investigation of cortical reactivity and connectivity elicited by TMS with high spatial specificity and temporal resolution (Bortoletto et al., 2015; Hernandez-Pavon, Veniero, et al., 2023; Ilmoniemi et al., 1997). For this feature, TMS-evoked potentials (TEPs) are increasingly explored as potential biomarkers of various neuropsychiatric disorders, including mood disorders, schizophrenia, neurodegenerative disorders, and epilepsy, in which connectivity alterations have been highlighted (e.g., Bagattini et al., 2019; Cao et al., 2021; S. Casarotto et al., 2019; Casula et al., 2023; Gefferie et al., 2023; Tremblay et al., 2019). Despite the growing use of TMS-EEG, the effect of stimulation parameters (e.g., coil orientation, technical features of TMS pulses) on the recorded signal remains ambiguous, possibly contributing to TEPs variability, thereby hindering their interpretation. To improve TMS-EEG application in both basic and clinical research, it is mandatory to understand which technical parameters contribute to shaping the recorded signal and how they can be manipulated to optimize signal measurement.

Among critical TMS parameters, there is evidence that pulse waveform and current direction are crucial for stimulation outcomes, determining the neural population preferentially activated by TMS in the target area. In the last three decades, TMS literature has used the motor system as the operative model for such investigation due to the easy read-out of the stimulation outcomes given by motor-evoked potentials (MEPs) which, in turn, can be used as proxies of M1 exogenous activation (e.g., Casula et al., 2018; Cirillo & Byblow, 2016; Corp et al., 2021; Davila-Pérez et al., 2018; Di Lazzaro et al., 2001; D’Ostilio et al., 2016; Kammer et al., 2001; Sakai et al., 1997; Sommer et al., 2006, 2018). Of interest for the present work, Aberra and colleagues (2020) recently developed a computational model for the human motor system showing that posterior-anterior (PA)-directed TMS current over M1 activates precentral gyrus neurons located more posteriorly compared to the ones activated with anterior-posterior (AP)-directed currents (Aberra et al., 2020). Notably, the caudal part of the precentral gyrus hosts cortico-motoneuronal cells, which reach spinal motoneurons through fast monosynaptic projections; by contrast, neurons in the rostral portion of this gyrus connect with spinal motoneurons through polysynaptic or slow monosynaptic connections (Siebner et al., 2022).

Importantly, it is possible, but still little explored, that pulse waveform and current direction determine which connections with other cortical regions and brain structures are preferentially activated. Studies on the animal model show that neuronal populations of the rostral and caudal parts of the precentral gyrus have direct connections with distinct nearby cortical regions. For instance, studies in monkeys found that neurons in the rostral portion of M1 are mainly connected to somatosensory and higher-order motor areas, while caudal populations communicate with the primary somatosensory cortex (Stepniewska et al., 1993); moreover, these neurons show slightly different cortico-thalamic connections (Matelli et al., 1989; Stepniewska et al., 1994). Furthermore, studies in humans employing repetitive and paired-pulse TMS suggest that the stimulation of M1 and interconnected areas lead to distinct neurophysiological aftereffects according to the TMS parameters exploited (e.g., Delvendahl et al., 2014; Federico & Perez, 2017; Hamada et al., 2014; Koch et al., 2013; Ni et al., 2011; Tings et al., 2005). Hence, we can hypothesize that different TMS parameters activate not only dissociable neurons of the corticospinal tract – as already thoroughly investigated (for reviews, see: Di Lazzaro et al., 2018; Spampinato, 2020) – but also divergent cortico-cortical pathways and networks, according to the M1 neural population primarily stimulated.

So far, a few studies have addressed how current direction and pulse waveform influence TMS-EEG outcomes after targeting M1 (e.g., Bonato et al., 2006; Casula et al., 2018; Guidali et al., 2023), finding modulations of signal amplitude and latency mainly in the early responses (i.e., <50 ms from TMS pulse). Interestingly, in the study conducted by our research group, we focused on an early M1-TEP component possibly reflecting transcallosal inhibition (i.e., M1-P15) and found that it was absent when coil orientation was changed in monophasic stimulation conditions (Guidali et al., 2023), suggesting that signal spread through the corpus callosum may depend on the TMS parameters exploited.

Critically, when extracting TEP waveforms, as in the studies mentioned above, only a portion of the available data is used, mainly focusing on the response’s amplitude of some target electrodes in a specific time window, neglecting whole-scalp signal temporal transitions. However, dissimilarities in TEP amplitudes and latencies cannot disentangle whether changing stimulation parameters affects the degree of activation of the same neural circuitry or the cortical pathways involved. Topographic analyses can overcome this issue, providing valuable insights into the spatiotemporal dynamics of cortical signal propagation (Michel et al., 2024; Vaughan, 1982). Indeed, differences in topographies reflect variations in the coordination of neuronal activity over time and the activation of distinct cortical sources and networks within the brain (Ding et al., 2024; Sulcova et al., 2022). Hence, topographic analysis of event-related potentials (ERPs) can detect the consistency of the neural activation elicited by TMS across subjects within the same experimental condition (test for topographic consistency; Habermann et al., 2018; Koenig et al., 2011) and can also be used to recognize fixed topographies characterising the data for specific time periods (i.e., microstates; Lehmann et al., 1987; Michel & Koenig, 2018; Tarailis et al., 2024). Importantly, these microstates have been shown to closely align with ERP components, thus providing information on typical component topography and duration that complements the classic analysis of peak amplitude and latency (Murray et al., 2008; Sulcova et al., 2022). In our case, microstate analysis can prove instrumental in comprehending whether alterations in stimulation parameters result in distinct brain functional states, inferring the activation of distinct cortical networks (e.g., Sulcova et al., 2022). This knowledge complements the information provided by waveforms analysis to better understand the role of TMS parameters in the recorded response.

Therefore, taking advantage of TMS-EEG, our study aims to investigate the impact of TMS current direction and pulse waveform on the activation of different cortico-cortical circuits after M1 stimulation, shedding light on possible sources of variability in TEP recording. To this end, we exploited three different current directions (PA, AP, and latero-medial – LM) and two pulse waveforms (monophasic and biphasic) for M1 stimulation while concurrently registering EEG activity. Data were analysed using two different but complementary approaches. Firstly, we assessed whether amplitudes and latencies of the classic TEP components elicited over M1 (i.e., N15, P30, N45, P60, N100, and P180; Beck et al., 2024) are influenced by stimulation parameters changes. Secondly, we investigated whether scalp topographies change among conditions and if microstates can be informative to better characterize TMS-induced spread of M1 activity and, in turn, the activation of distinct circuitries when stimulation parameters are modulated.

## 2. MATERIALS AND METHODS

### 2.1. Participants

The dataset used in this study was taken from the original work by Guidali et al. (2023), and raw data can be found at https://gin.g-node.org/Giacomo_Guidali/Guidali_et_al_2023_EJN_RR. It comprised 40 right-handed healthy participants ranging in age from 18 to 50 years and meeting the criteria for TMS safety (Rossi et al., 2021). Of the forty participants tested, we excluded those who required a stimulation intensity exceeding 90% of the maximal stimulator output (six participants) and those who did not complete all experimental blocks (two participants). Therefore, the final analysed sample in this experiment comprises 32 participants [18 females, median age: 26 years (range: 20-49 years); median education: 16 years (range: 13-21 years); median Edinburgh (Oldfield, 1971) score: 83% (range: 42%- 100%)]. The research was conducted at the Neurophysiology Laboratory of the IRCCS Istituto Centro San Giovanni di Dio Fatebenefratelli in Brescia, Italy. This study followed the ethical guidelines outlined in the Declaration of Helsinki and received approval from the local ethics committee at IRCCS Istituto Centro San Giovanni di Dio Fatebenefratelli (reference number: 102-2021). Datasets and statistical analyses for the present work can be found on Open Science Framework – OSF at: https://osf.io/8wrgm/?view_only=5ecb841ab7b645f9821486625978cc9c.

### 2.2. Experimental procedure

Participants took part in a single-session experiment composed of seven blocks of TMS-EEG co-registration (see Guidali et al., 2023 for more details). Each block was characterized by a combination of current direction (PA, AP, LM) and pulse waveform (monophasic or biphasic). It must be noted that for biphasic stimulation, we always refer to the second phase of the TMS pulse (e.g., biphasic AP-PA stimuli are denoted as biphasic PA). Each experimental block included a consecutive evaluation of the rMT, a TMS-electromyography (EMG), and a TMS-EEG recording. TMS-EMG and the seventh TMS-EEG block in which one hand was contracted have not been further analysed in the present work. MEP patterns taken from our original study (Guidali et al., 2023) are reported in the **Supplemental Figure 1**. rMT was measured through the best parameter estimation by sequential testing (PEST) procedure (Awiszus, 2003). TMS-EEG recordings consisted of 80 TMS pulses, with an inter-pulse interval jittered between 4000 and 6000 ms. TMS intensity was adjusted to 110% of the participant’s rMT of each block. rMT values in the six conditions were reported in the **Supplemental Figure 2.** Block order was counterbalanced between participants following a Latin square design (**Figure 1**).

**Figure 1.**
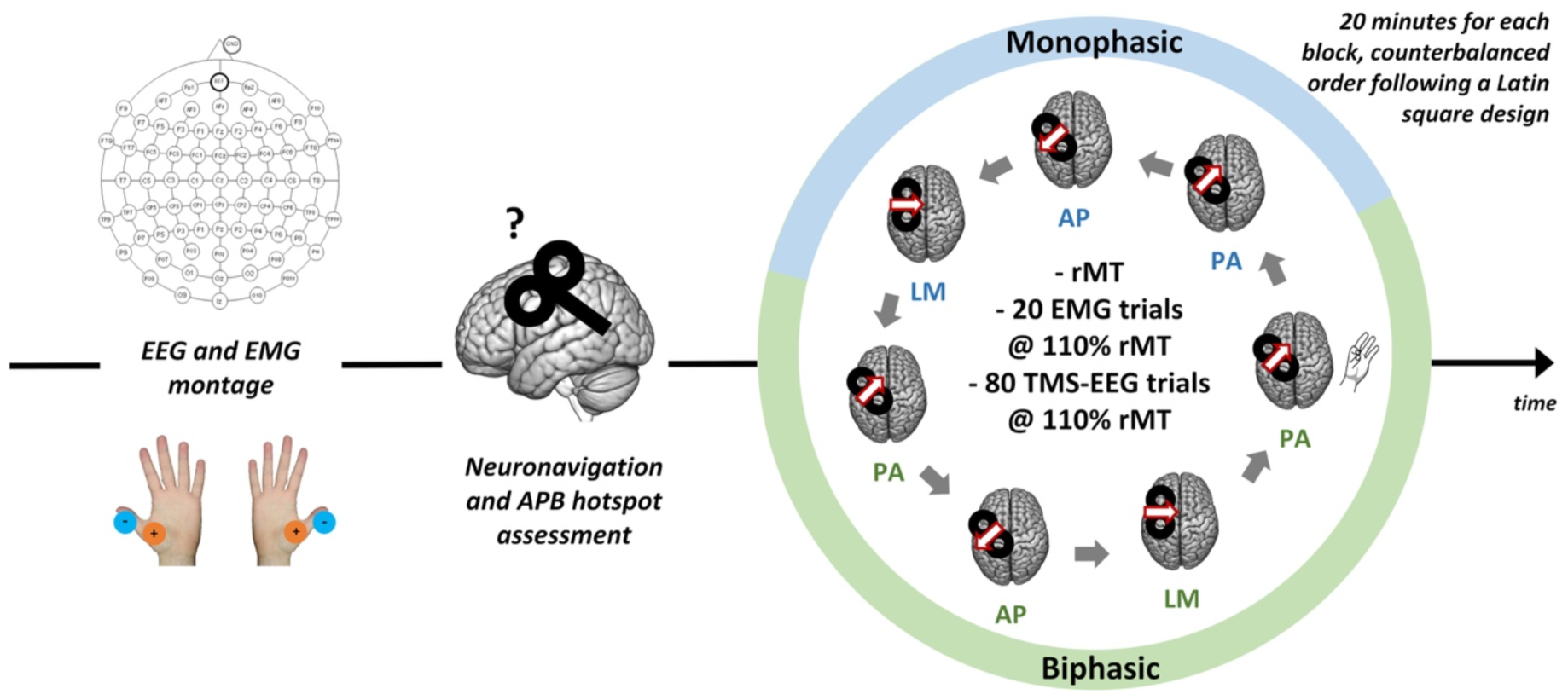
Experimental procedure of the original study by Guidali et al. (2023). Participants underwent seven stimulation blocks after EEG and EMG montage, neuronavigation, and APB hotspot assessment. Each block, characterized by the combination of a specific pulse waveform and current direction, was composed of rMT computation, 20 trials recording MEPs, and 80 trials recording TEPs. For the aim of the present work, the block in which one hand was contracted has not been analysed. In all our figures, red and white arrows over TMS coils represent the direction of the induced current in M1.

Throughout the experiment, participants were seated in a dimly lit room with their forearms comfortably resting on a table, positioned in front of a computer screen showing a fixation cross. Before starting with the experimental blocks, the EEG cap and EMG electrodes over the right *abductor pollicis brevis* (APB) were applied. Then, the motor hotspot for the right APB was determined using a biphasic stimulator and a PA current direction. This hotspot was selected as the consistent stimulation target during the experiment.

The participants wore noise-cancelling earphones, and white noise was played during blocks to minimize the contamination of TMS-locked auditory artifacts in the recorded EEG trace (Biabani et al., 2019). The volume of the white noise was individually adjusted to a maximum of 90 dB. A thin layer of foam was placed beneath the TMS coil to attenuate the sensory stimulation caused by TMS.

### 2.3. TMS-EEG recording

A figure-of-eight coil (Magstim model Alpha B.I. Coil Range, diameter: 70 mm) was used for all stimulation blocks. For monophasic blocks, a Magstim 200^2^ stimulator was used; for biphasic blocks, we employed a Magstim Rapid^2^ stimulator (Magstim, Whitland, UK). The current direction was changed depending on the block. For PA blocks, the coil was oriented 45° to the midline. For AP blocks, the coil was oriented 180° to the PA orientation. For LM blocks, the coil was oriented 90° to the midline. Neuronavigation procedures were performed using SofTaxic Optic 3.4 neuronavigation software (EMS, Bologna, Italy; www.softaxic.com) to check the TMS coil position continuously.

EEG recording was carried out using an EEG system compatible with TMS (g.HIamp multichannel amplifier, g.tec medical engineering GmbH). EEG data was collected from 74 electrodes (EasyCap, Brain Products GmbH, Munich, Germany) placed on the scalp following the 10–10 international system, referenced to FPz, and with the ground electrode on the tip of the nose. Data were acquired with a sampling rate of 9.6 kHz, and skin-electrode impedance was consistently maintained below 5 kΩ. The EEG signal was also visually monitored before and during the recordings to identify and address any apparent artifacts, such as prolonged decays, noisy channels, and line noise.

### 2.4. EEG preprocessing and TEP extraction

We used the same pre-processed TEP data of Guidali et al. (2023). The pre-processing pipeline, implemented in MATLAB R2020b, combining EEGLAB v.2020.0 (Delorme & Makeig, 2004) and Fieldtrip v.20190905 functions (Oostenveld et al., 2011), is available at https://gin.g-node.org/Giacomo_Guidali/Guidali_et_al_2023_EJN_RR/src/master/Script%20preprocessing%20EMG-EEG/preprocessingTEP_pipeline.m and it is the pipeline commonly used by our research group (e.g., Guidali et al., 2023; Zazio et al., 2022, 2024). The pipeline comprised the source-estimate-utilizing noise-discarding (SOUND) algorithm (Mutanen et al., 2018) to clean bad channels and reduced measurement noise, independent component analysis (ICA) to selectively remove ocular artifacts, and the signal-space projection and source-informed reconstruction (SSP-SIR) algorithm (Mutanen et al., 2016) to clean TMS-related muscular artifact. For further details, and the description of all preprocessing steps, see: Guidali et al., 2023.

To compute M1-TEP peak amplitude and latency in the experimental conditions of interest, we individuated six well-established TEP components generated through M1 stimulation: N15, P30, N45, P60, N100, and P180 (Beck et al., 2024; Farzan & Bortoletto, 2022). The P15 peak (Bortoletto et al., 2021; Zazio et al., 2022) was already analysed in our original work (Guidali et al., 2023) and not further considered here. For each of these peaks, we performed the following steps. First, we examined the existing literature to identify the typical time window within which each TEP component is generally measured (e.g., Belardinelli et al., 2021; Farzan et al., 2013; Gordon et al., 2021; Rogasch & Fitzgerald, 2013; Zazio et al., 2021). Afterward, we verified the existence of a signal deflection within this temporal window in our grand grand-average (i.e., collapsing all conditions), and we re-defined the time interval so that it did not overlap with the ones chosen for the neighbouring TEP peaks. The signal was then averaged from the four electrodes corresponding to the peak location of each TEP component (similar to Guidali et al., 2023). Then, its amplitude and latency were computed for all participants in each experimental condition. The chosen time windows and electrodes for each M1-TEP component are reported in **Table 1**.

**Table 1.** Time of interest (TOI) and region of interest (ROI) for the six extracted M1-TEP components.

| *TEP component* | *TOI (ms)* | *ROI (channels)* |
| --- | --- | --- |
| **N15** | 7-25 | C3, CP3, CP1, CP5 |
| **P30** | 25 - 38 | Cz, FCz, C1, FC1 |
| **N45** | 38 - 55 | F4, FC4, FC2, F2 |
| **P60** | 55 - 75 | C3, CP3, CP1, CP5 |
| **N100** | 90 - 130 | FCz, C1, FC1, FC3 |
| **P180** | 170 - 220 | F2, FCz, FC2, C2 |

### 2.5. Microstate analysis

To perform microstate analysis, we used the MATLAB-based toolbox Randomization Graphical User software (RAGU - Koenig et al., 2011), following the guidelines published in previous literature (Habermann et al., 2018; Koenig et al., 2014). First, EEG traces were downsampled to 960 Hz. Then, after loading the data of the six conditions of interest (PAmonophasic, APmonophasic, LMmonophasic, PAbiphasic, APbiphasic, and LMbiphasic) we detected five outliers using the algorithm proposed by the software – based on the Mahalanobis distance between subjects (Wilks, 1961). These subjects were then excluded from the following analysis. We defined the within-subject experimental design for two factors (current direction, pulse waveform) and set the randomization to 5000 runs with a significance level of p = .05. Before analysis, the subject-average data was normalized by its global field power (GFP). This permits the detection of differences in the spatial distribution of the sources at the scalp level while minimizing the differences due to the variability in TEP amplitude (Habermann et al., 2018). Finally, we tested topographic consistency and microstate analysis in TEP data between 5 and 400 ms after stimulation. The GFP of the grand average TEP in each condition was compared to the null distribution to perform the test for topographic consistency. This distribution was created by shuffling the channel positions of individual-average TEPs (resulting in random topographies) and recomputing the mean GFP 5000 times (Habermann et al., 2018).

To identify the optimal number of maps for microstate analysis, we used the cross-validation approach with 50 iterations, as introduced by Koenig and colleagues (Koenig et al., 2014). In each step, the dataset was randomly divided into a learning set, used to build microstate models with number of classes increasing from 3 to 12, and a test set, used to verify the created classes and establish the amount of variance explained by each model (for more information on the procedure, see Koenig et al., 2014; Sulcova et al. 2022). The cross-validation results suggested that our data can be optimally explained by 6 microstate classes, as the 6-map model was the first one reaching the plateau value of global explained variance. Therefore, we used the k-means clustering algorithm with 250 iterations to segment the TEPs into microstates defined by 6 different topographic maps. In each iteration of this process, the group-average data of all conditions was concatenated, 6 scalp topographies were randomly selected from this dataset, and the topography most correlated with each datapoint was used to label that datapoint. The templates were then refined by averaging the signal across the datapoints labelled with the same map. Finally, the set of template maps that showed the highest global explained variance was retained.

### 2.6. Statistical analysis

Statistical significance was set in all our analysis at p <.05. For repeated-measures analyses of variance (rmANOVAs) and analyses of covariance (ANCOVAs), data analysis was performed using Jamovi software (Version 2.3.21; The Jamovi Project, 2023), while robust ANOVAs and generalized linear mixed models were conducted with R Studio (Version 1.2.5019 R Core Team, 2020) using ‘WRS 2’ (Mair & Wilcox, 2020) and ‘lme4’ (Bates et al., 2015) packages.

#### 2.6.1. TEPs

For each TEP component, participants showing amplitude and/or latency values exceeding ± 2.5 SD from the group’s mean were considered outliers and excluded from the analysis (range 3-5; see **Supplemental Table 1** for more information). We analysed latency and amplitude of each TEP peak with separate rmANOVAs with factors ‘Pulse waveform’ (monophasic, biphasic) and ‘Current direction’ (PA, AP, LM). Then, to verify the possible contribution of TMS intensity on peak amplitude modulation, we conducted ANCOVAs for each pulse waveform, with the factor ‘Current direction’ (PA, AP, LM), and covarying the TMS intensity value used for the corresponding block. Data sphericity was assessed with Mauchly’s test, and if not confirmed, the Greenhouse–Geisser correction was used. For post-hoc tests, Tukey correction for multiple comparisons was applied. Partial eta-squared (ƞp^2^) for rmANOVA and Cohen’s *d* for t-test were reported as effect size values.

The normality of data distribution was checked for each variable and, if needed, (*a*) square root, (*b*) base-ten logarithm, and (*c*) inverse transformations were conducted to identify the transformation making data distribution closer to normality (same procedure as in Guidali et al., 2023). The P60 amplitude distribution was normalized with a base-ten logarithm transformation. For the latency of the P30, N45, P60, N100, and P180 components, none of these transformations worked; therefore, two (one for each pulse waveform - monophasic and biphasic) robust one-way rmANOVAs based on trimmed means (20% trimming level) with three factors (i.e., AP, PA, LM) were performed on the raw data (Mair & Wilcox, 2020).

#### 2.6.2. Microstates

As a preliminary analysis, we run the test for topographic consistency to verify the presence of consistent topographies across conditions. Then, to specifically investigate whether TMS parameters modulated microstate properties, we analysed, for every class individuated from microstate extraction (see **2.5**), (i) the area under the curve (AUC) – i.e., the sum of the GFP amplitude for each timepoint in which they are detected – and (ii) their duration – i.e., the sum of all timepoints in which they occurred – with a series of ‘Pulse waveform’ X ‘ Current direction’ rmANOVAs, as for TEP indexes (see previous paragraph). Microstate duration showed a normal distribution, while none of our planned transformations made microstate AUC data closer to normality. Therefore, for this variable, we proceeded with a series of robust rmANOVAs (Mair & Wilcox, 2020) as described above. Furthermore, we specifically explored whether different stimulations activated distinct circuits. To this aim, we examined changes in the spatiotemporal pattern of responses by looking at the order of microstate appearance across conditions. Considering that different topographies represent the activation of different cortical sources, changes in the sequence of transitions from one microstate to another likely reflect different patterns of effective connectivity. Therefore, we analysed microstate onset values, i.e., the first timepoint at which a microstate class appeared, adopting a generalized linear mixed model (gamma distribution, identity link). ‘Microstate class’ (1, 2, 3, 4, 5, and 6), ‘Stimulation condition’ (monophasic PA, monophasic AP, monophasic LM, biphasic PA, biphasic AP, and biphasic LM), and their interaction were included as fixed effects. Subjects were included as the random effect. Post-hoc tests were corrected for multiple comparisons using the Benjamini-Hochberg method. Given the aim of this analysis, in the exploration of post-hoc comparisons, we considered only contrasts between microstate classes within the same stimulation condition, with particular attention to cases where the onset order of two classes swapped between stimulation conditions.

## 3. RESULTS

### 3.1. TEP peaks

TEP grand-averages and components’ topographical maps for each stimulation condition are depicted in **Figure 2**. The temporal succession of M1-TEP peaks and related topographies align with the ones reported by previous TMS-EEG literature stimulating the motor cortex (for reviews, see: Beck et al., 2024; Farzan & Bortoletto, 2022). Visually striking differences among our experimental conditions can be specifically detected in the first 50 ms from the TMS pulse: monophasic waveforms show different patterns of topographies among current directions, with AP currents presenting the greatest early components.

**Figure 2.**
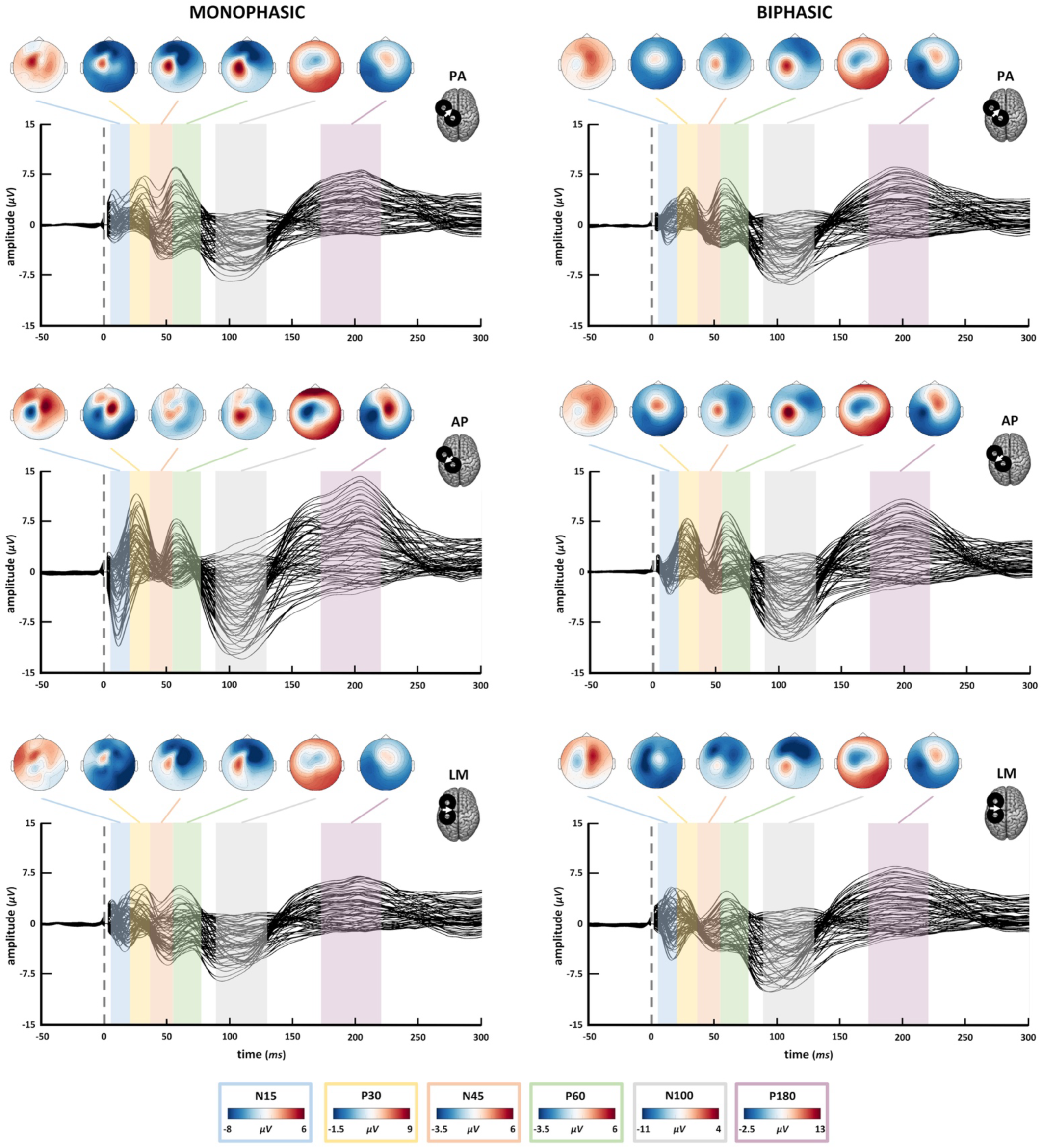
TEP grand averages for all conditions and topographical maps of all components. For the specific time intervals of each component, see Table 1. The voltage scale used for topographies changes across components, see the legend at the bottom for the corresponding scale.

Means (M) and standard errors (SE) of M1-TEP components’ amplitude and latency in the six experimental conditions are reported in **Table 2**.

**Table 2.** Amplitude and latency’s Ms and SEs of M1-TEP components in the six experimental conditions.

| *Variable* | *Pulse waveform* | *Current direction* | *N15*<br>*(M ± SE)* | *P30*<br>*(M ± SE)* | *N45*<br>*(M ± SE)* | *P60*<br>*(M ± SE)* | *N100*<br>*(M ± SE)* | *P180*<br>*(M ± SE)* |
| --- | --- | --- | --- | --- | --- | --- | --- | --- |
| *Amplitude*<br>*(μV)* | **Monophasic** | **PA** | 1.10 ± .5 | 6.65 ± .8 | -5.89 ± .5 | 6.76 ± .91 | -7.21 ± .8 | 7.85 ± .9 |
|  |  | **AP** | -8.34 ± .9 | 4.73 ± .9 | -0.84 ± .7 | 6.27 ± 1.16 | -12.77 ± 1.1 | 13.11 ± 1.1 |
|  |  | **LM** | -2.07 ± .8 | 5.44 ± .5 | -5.99 ± 0.4 | 4.24 ± .84 | -6.95 ± .8 | 6.7 ± .6 |
|  | **Biphasic** | **PA** | -2.13 ± .5 | 5.03 ± .5 | -3.8 ± .5 | 5.49 ± .9 | -9.17 ± .7 | 8.73 ± .7 |
|  |  | **AP** | -2.60 ± .5 | 6.89 ± .6 | -4.15 ± .5 | 7.21 ± 1.04 | -10.19 ± 1.1 | 9.86 ± .6 |
|  |  | **LM** | -3.97 ± .7 | 2.24 ± .7 | -3.95 ± .5 | 3.76 ± .96 | -10.08 ± .8 | 8.4 ± .7 |
| *Latency*<br>*(ms)* | **Monophasic** | **PA** | 12.36 ± .6 | 31.81 ± .7 | 48.42 ± .9 | 58.16 ± .7 | 106.14 ± 2.2 | 199.12 ± 2.5 |
|  |  | **AP** | 18.07 ± .9 | 29.92 ± .6 | 47.68 ± 1 | 57.35 ± .6 | 105.27 ± 2.3 | 193.47 ± 3 |
|  |  | **LM** | 17.87 ± .9 | 31.3 ± .8 | 48.37 ± 1 | 61.72 ± 1.2 | 104.58 ± 2.1 | 195.11 ± 3.4 |
|  | **Biphasic** | **PA** | 14.33 ± .8 | 30.27 ± .7 | 49.02 ± .9 | 58.54 ± .6 | 105.39 ± 2.2 | 195.64 ± 2.6 |
|  |  | **AP** | 13.64 ± .8 | 30.66 ± .6 | 49.45 ± .8 | 58.11 ± .6 | 103.02 ± 1.8 | 194.3 ± 3.1 |
|  |  | **LM** | 15.69 ± .9 | 31.95 ± .8 | 50.12 ± 1 | 60.56 ± 1.1 | 101.51 ± 1.7 | 194.45 ± 3 |

#### 3.1.1. Amplitude

Overall, all six TEP peak amplitudes were significantly modulated by stimulation parameters with an interaction effect of ‘Current direction’ X ‘Pulse waveform’ (for all rmANOVAs main effects and interactions, see **Supplemental Table 2)**.

Specifically, post-hoc comparisons for the N15 (*F*2,56 = 26.53, *p* < .001, ƞp^2^ = .487) showed that monophasic AP led to the greatest (i.e., more negative) amplitude and the monophasic PA direction led to the smallest N15. The latter condition was characterised by the absence of a negative component in the analysed electrodes (for significant post-hoc comparisons, see: **Table 3**; **Figure 3a**).

**Figure 3.**
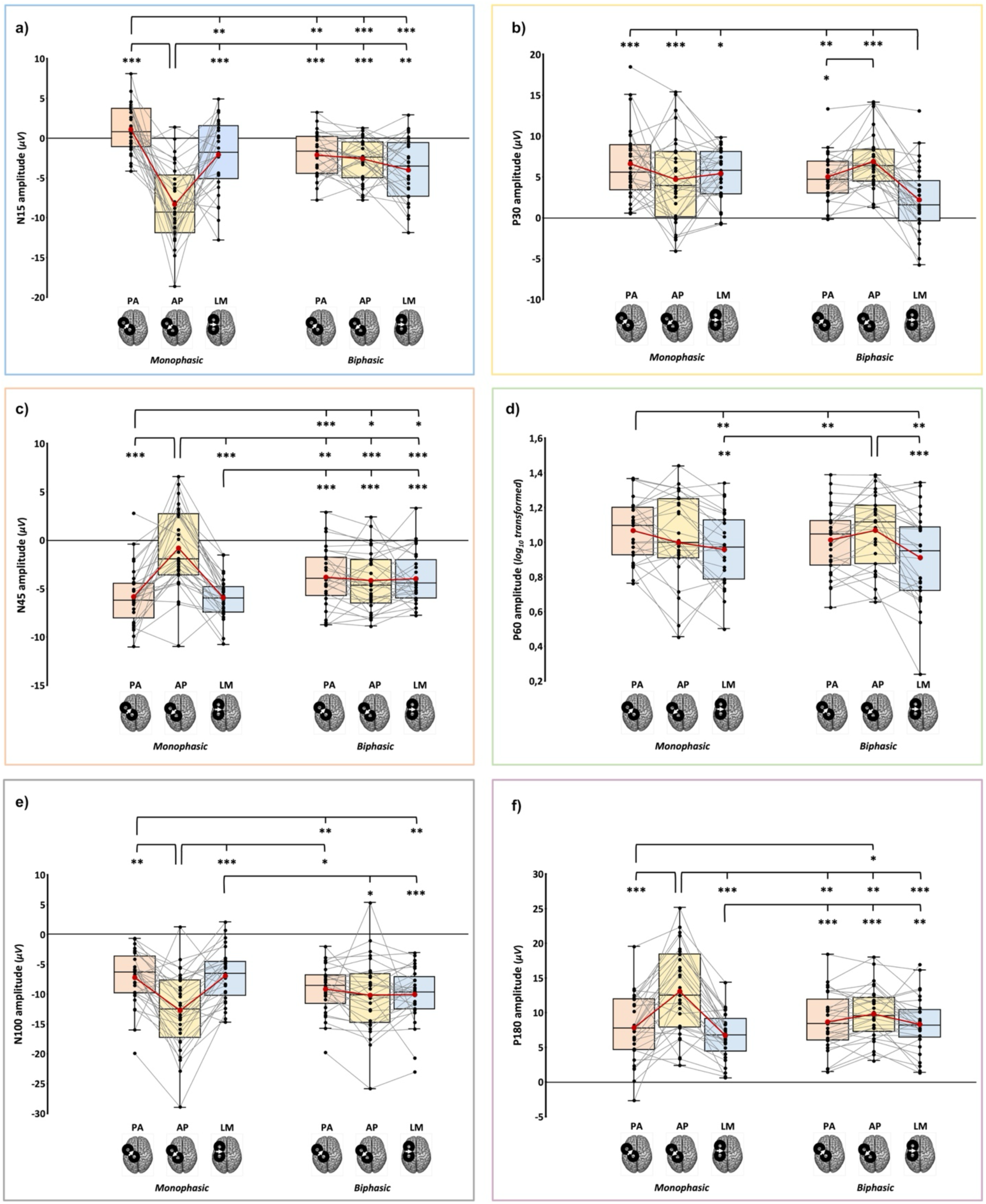
Amplitude of N15 (**a**), P30 (**b**), N45 (**c**), P60 (**d**), N100 (**e**), and P180 (**f**) components in the six experimental blocks. In the box-and-whiskers plots, red dots and lines represent the means of the distributions. The centre line depicts their median values. Black dots and grey lines show individual scores. The box contains the 25th to 75th percentiles of the dataset. Whiskers extend to the largest observation, which falls within the 1.5 times interquartile range from the first/third quartile; significant p-values of corrected post hoc comparisons are reported (* = p <.05; ** = p <.01; *** = p <.001).

**Table 3.** Significant post-hoc comparisons (Tukey corrected) for TEP peaks’ amplitude.

| *TEP component* | *Condition* | *versus* | *t* | *p<sub>Tukey</sub>* | *Cohen's d* |
| --- | --- | --- | --- | --- | --- |
| **N15** | **AP<sub>monophasic</sub>** | **PA<sub>monophasic</sub>** | -7.9 | <.001 | -1.47 |
|  |  | **LM<sub>monophasic</sub>** | -4.7 | <.001 | -0.87 |
|  |  | **PA<sub>biphasic</sub>** | -7.1 | <.001 | -1.32 |
|  |  | **AP<sub>biphasic</sub>** | -5.6 | <.001 | -1.04 |
|  |  | **LM<sub>biphasic</sub>** | -3.9 | .007 | -0.72 |
|  | **PA<sub>monophasic</sub>** | **AP<sub>monophasic</sub>** | 7.9 | <.001 | 1.47 |
|  |  | **LM<sub>monophasic</sub>** | 3.8 | .008 | 0.71 |
|  |  | **PA<sub>biphasic</sub>** | 4.6 | .001 | 0.85 |
|  |  | **AP<sub>biphasic</sub>** | 6.1 | <.001 | 1.13 |
|  |  | **LM<sub>biphasic</sub>** | 7.7 | <.001 | 1.43 |
| **P30** | **LM<sub>biphasic</sub>** | **PA<sub>monophasic</sub>** | -5 | <.001 | -0.93 |
|  |  | **AP<sub>monophasic</sub>** | -3.5 | .02 | -0.64 |
|  |  | **LM<sub>monophasic</sub>** | -4.6 | <.001 | -0.86 |
|  |  | **PA<sub>biphasic</sub>** | -4.4 | .002 | -0.82 |
|  |  | **AP<sub>biphasic</sub>** | -6 | <.001 | -1.12 |
|  | **PA<sub>biphasic</sub>** | **AP<sub>biphasic</sub>** | -3.39 | .023 | -.63 |
| **N45** | **AP<sub>monophasic</sub>** | **PA<sub>monophasic</sub>** | 6.3 | <.001 | 1.17 |
|  |  | **LM<sub>monophasic</sub>** | 8.4 | <.001 | 1.55 |
|  |  | **PA<sub>biphasic</sub>** | 4.1 | .004 | .77 |
|  |  | **AP<sub>biphasic</sub>** | 5.1 | <.001 | .95 |
|  |  | **LM<sub>biphasic</sub>** | 5.1 | <.001 | .94 |
|  | **PA<sub>monophasic</sub>** | **PA<sub>biphasic</sub>** | -5.5 | <.001 | -1.03 |
|  |  | **AP<sub>biphasic</sub>** | -3.6 | .014 | -.67 |
|  |  | **LM<sub>biphasic</sub>** | -3.7 | .012 | -.68 |
|  | **LM<sub>monophasic</sub>** | **PA<sub>biphasic</sub>** | -5.5 | <.001 | -1.02 |
|  |  | **AP<sub>biphasic</sub>** | -4 | .005 | -.74 |
|  |  | **LM<sub>biphasic</sub>** | -5.6 | <.001 | -1.04 |
| **P60** | **PA<sub>monophasic</sub>** | **PA<sub>biphasic</sub>** | 3.9 | .008 | .75 |
|  | **LM<sub>monophasic</sub>** | **PA<sub>monophasic</sub>** | 4.1 | .004 | .8 |
|  |  | **AP<sub>biphasic</sub>** | 4.1 | .004 | .79 |
|  | **LM<sub>biphasic</sub>** | **PA<sub>monophasic</sub>** | 4.1 | .004 | .79 |
|  |  | **AP<sub>biphasic</sub>** | 4.7 | <.001 | .91 |
| **N100** | **AP<sub>monophasic</sub>** | **PA<sub>monophasic</sub>** | -4.59 | .001 | -.85 |
|  |  | **LM<sub>monophasic</sub>** | -5.83 | <.001 | -1.09 |
|  |  | **PA<sub>biphasic</sub>** | 3.1 | .044 | -.58 |
|  | **PA<sub>monophasic</sub>** | **PA<sub>biphasic</sub>** | 3.8 | .008 | .71 |
|  |  | **LM<sub>biphasic</sub>** | -4 | .005 | -.74 |
|  | **LM<sub>monophasic</sub>** | **AP<sub>biphasic</sub>** | -3.3 | .025 | -.62 |
|  |  | **LM<sub>biphasic</sub>** | 5.3 | <.001 | .98 |
| **P180** | **AP<sub>monophasic</sub>** | **PA<sub>monophasic</sub>** | 5.1 | <.001 | .93 |
|  |  | **LM<sub>monophasic</sub>** | 7.1 | <.001 | 1.34 |
|  |  | **PA<sub>biphasic</sub>** | 5.3 | <.001 | .99 |
|  |  | **AP<sub>biphasic</sub>** | 5.3 | <.001 | .79 |
|  |  | **LM<sub>biphasic</sub>** | 4.5 | .001 | .84 |
|  | **LM<sub>monophasic</sub>** | **PA<sub>biphasic</sub>** | -5.2 | <.001 | -.97 |
|  |  | **AP<sub>biphasic</sub>** | -7.1 | <.001 | -1.32 |
|  |  | **LM<sub>biphasic</sub>** | -4.5 | .002 | -.83 |
|  | **PA<sub>monophasic</sub>** | **AP<sub>biphasic</sub>** | 4.2 | .003 | -.62 |

Regarding P30 (*F*1.65,46.11 = 11.37, *p* < .001, ƞp^2^ = .289), post-hoc tests showed that biphasic LM currents led to the smallest P30 component, which was significantly different from all the other stimulation conditions. Furthermore, among biphasic pulse waveforms, the PA direction evoked a P30 component with a lower amplitude than the AP one (**Figure 3b**).

For the N45 (*F*1.59,44.54 = 35.98, *p* < .001, ƞp^2^ = .562), post-hoc comparisons showed that monophasic pulses with AP current direction evoked the less negative N45 amplitude among all experimental conditions, and the analysis of single-subjects distribution (**Figure 3c**) suggests a great variability of N45 polarity in this condition. Conversely, the other two monophasic waveforms (i.e., PAmonophasic and LMmonophasic) led to more negative N45 than all three biphasic conditions (**Figure 3c**).

Considering the P60 (*F*1.32,34.4 = 5.286, *p* = .008, ƞp^2^ = .168), the monophasic PA current direction evoked a higher amplitude than the homolog biphasic direction. Furthermore, the two LM directions led to significantly smaller P60 than PAmonophasic and APbiphasic (**Figure 3d**).

For the N100 (*F*2,56 = 22.06, *p* < .001, ƞp^2^ = .441), monophasic AP direction evoked components with more negative amplitudes than the other two monophasic conditions and biphasic PA currents. Monophasic PA, instead, evoked fewer negative components than the homolog and LM biphasic directions. Monophasic LM showed a similar pattern to the latter – i.e., less negative N100 concerning the homolog and the AP biphasic directions (**Figure 3e**).

Finally, considering the P180 (*F*1.56,43.65 = 19.7, *p* < .001, ƞp^2^ = .413), post-hoc comparisons showed that monophasic AP elicited the highest P180 with respect to all other five experimental conditions. In turn, the monophasic LM current direction generated lower amplitudes than the three biphasic waveforms, and monophasic PA currents led to a smaller P180 than the one recorded using the biphasic AP direction (**Figure 3f**).

Finally, ANCOVAs - run to verify the possible contribution of stimulation intensity on peak amplitude modulation - showed a significant effect of TMS intensity only for the P180 component for both monophasic (*F*1,83 = 14.97, *p* < .001, ƞp^2^ = .153) and biphasic waveforms (*F*1,83 = 9.33, *p* = .003, ƞp^2^ = .101). Interestingly, for biphasic stimulation, no significant effect of ‘Current direction’ at the net of TMS intensity was found (*F*2,83 = 1.88, *p* = .16, ƞp^2^ = .043), suggesting that the differences in stimulator’s intensity mainly drove amplitude modulations found in biphasic conditions. Conversely, for monophasic pulse waveform, the factors ‘Current direction’ (*F*2,83 =3.94, *p* = .023, ƞp^2^ = .087) still showed a significant effect, and post-hoc comparisons showed a significantly higher amplitude evoked by AP compared to LM (*t83* = 2.8, *p* = .019, *d* = 1.34), as already highlighted by the main analysis.

For the other five components, the contribution of TMS intensity was never statistically significant (all *F*s < 3.12, all *p*s > .081).

#### 3.1.2. Latency

Considering latency, the N15 and P60 were the only components showing significant modulations (for all rmANOVAs main effects and interactions, see **Supplemental Table 3**).

The N15 was significantly modulated, presenting an interaction effect ‘Current direction’ X ‘Pulse waveform’ (*F*2,56 = 10.39, *p* < .001, ƞp^2^ = .271). Post-hoc comparisons revealed that monophasic AP currents led to an N15 component with a significantly shorter latency than the other two monophasic conditions (vs. PAmonophasic: *t* = -5.4, *p* < .001, *d* = -.98; vs. LMmonophasic: *t* = -4.4, *p* = .002, *d* = -.81) and the biphasic LM direction (*t* = -3.4, *p* = .022, *d* = -.63). In turn, monophasic PA and LM currents led to higher latency values compared to biphasic PA (vs. PAmonophasic: *t* = -4.4, *p* = .002, *d* = -.81; vs. LMmonophasic: *t* = -3.4, *p* = .023, *d* = -.63) and AP conditions (vs. PAmonophasic: *t* = -3.6, *p* = .013, *d* = -.67; vs. LMmonophasic: *t* = -3.1, *p* = .047, *d* = -.57; **Figure 4a**).

**Figure 4.**
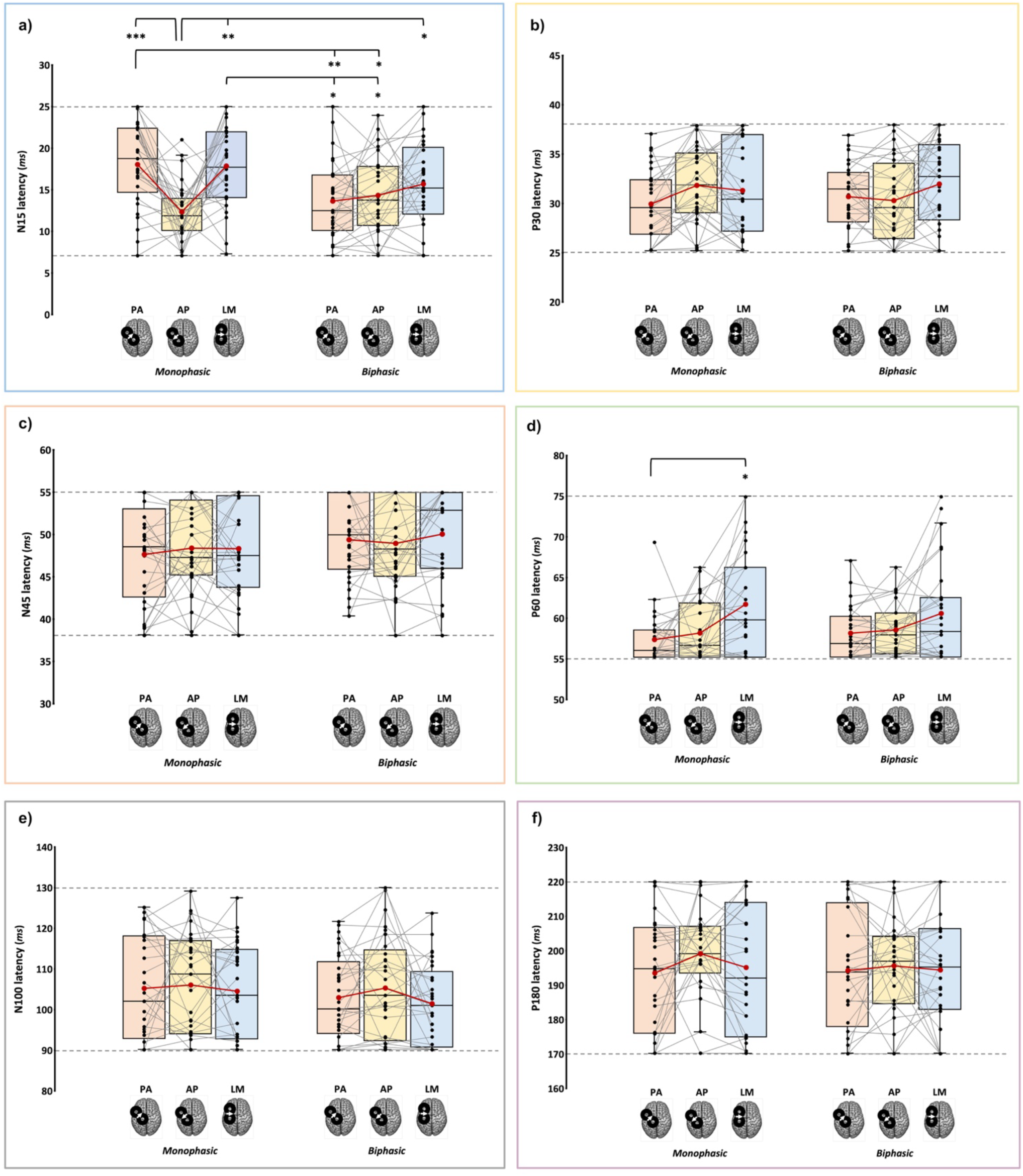
Latency of N15 (**a**), P30 (**b**), N45 (**c**), P60 (**d**), N100 (**e**), and P180 (**f**) components in the six experimental blocks. In the box-and-whiskers plots, red dots and lines represent the means of the distributions. The centre line depicts their median values. Black dots and grey lines show individual scores. The box contains the 25th to 75th percentiles of the dataset. Whiskers extend to the largest observation, which falls within the 1.5 times interquartile range from the first/third quartile; significant p-values of corrected post hoc comparisons are reported (* = p <.05; ** = p <.01; *** = p <.001).

Moreover, P60 latencies recorded with monophasic pulses were modulated by current direction (*F*1.65,26.48 = 5.57, *p* =.013). Indeed, LM currents evoked P60 with a significantly longer latency than PA ones (*ψ* =.003, *p* =.012; **Figure 4d**).

The other four components did not differ in latencies according to the current direction and the pulse waveform exploited (all *F*s < 2.56, all *p*s >.092; **Figure 4**).

### 3.2. Microstate analysis

Results of the test for topographic consistency showed that statistical significance was reached at every time point; therefore, a high consistency characterized all conditions, and the whole epoch (0-400 ms) was considered in the following microstates analysis (**Supplemental Figure 3**).

The adopted 6 microstate classes model explained a total variance of 91.34%. Visual inspection of microstate sequences shows that all six microstates are present in each stimulation condition. However, their succession and appearance differed across conditions, mainly in the first 50 ms after TMS. At later latencies, only the monophasic AP condition showed a distinct sequence of microstates compared to the other conditions, with the absence of a microstate map showing predominant positivity over fronto-central electrodes (i.e., class 5). Overall, the monophasic AP condition showed the most distinct pattern compared to the other conditions, displaying seven microstates within the first 100 ms. Notably, between 100 and 250 ms post-TMS, i.e., during the N100 and P180 time windows, it showed longer periods of negative signal over left sensorimotor electrodes (class 6) and positive signal over right frontal electrodes (class 3; **Figure 5**).

**Figure 5.**
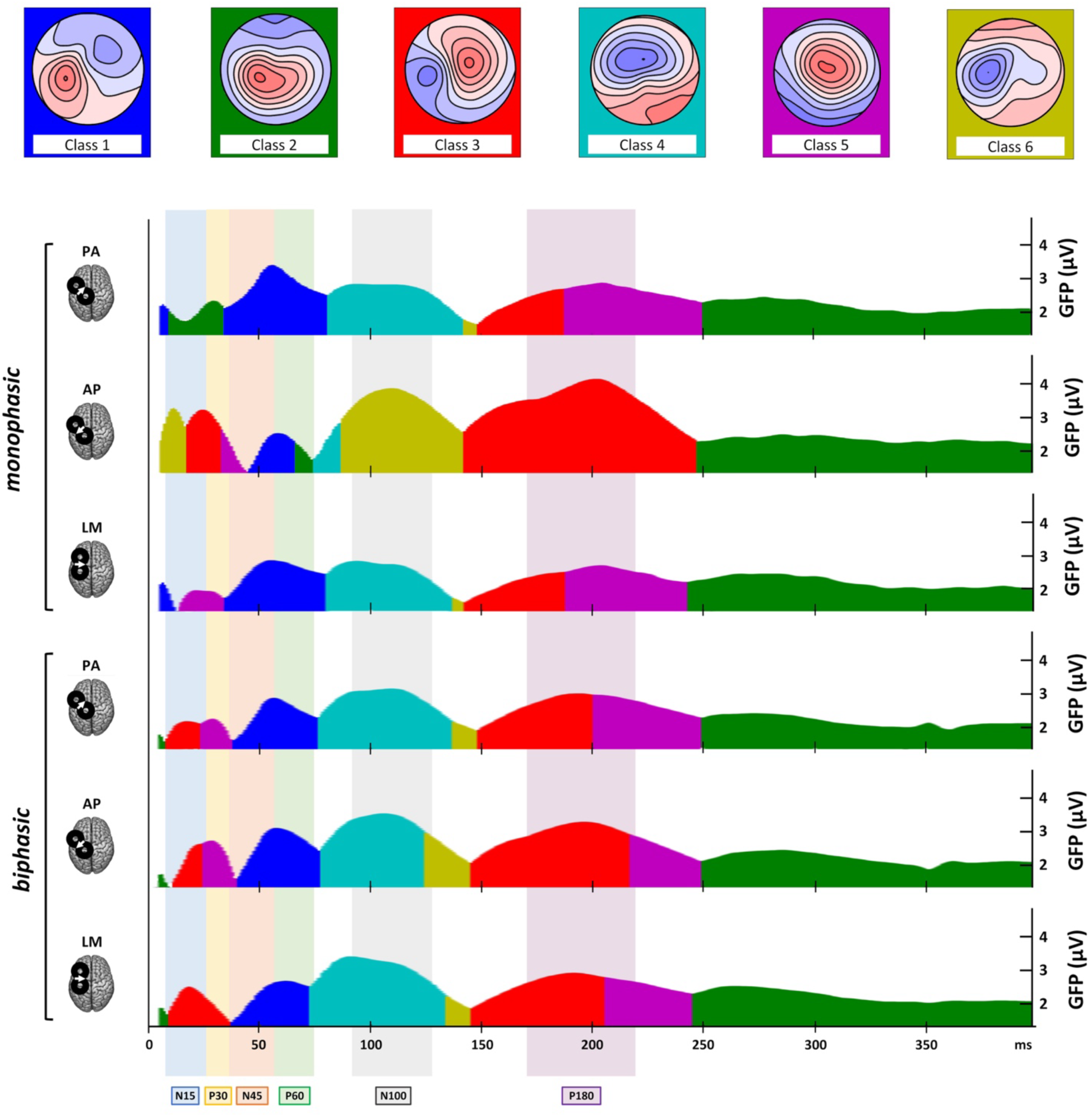
Microstate analysis results. Upper row: template maps of the 6 identified microstate classes. Main section: for all stimulation conditions, the proportion of the signal variance that can be explained by the model is coloured according to the microstate class dominating each time interval. Time windows attributed to specific TEP peaks are shaded in the colour referred to the corresponding component.

Microstates’ parameters analysis allowed us to define better and quantitatively characterize these patterns. The M and SE of the six microstates’ AUC, duration, and onset in the different experimental conditions are reported in **Table 4**.

**Table 4.** Microstate AUC, duration, and onset’s Ms and SEs of the six experimental conditions for the extracted classes.

| *Variable* | *Pulse waveform* | *Current direction* | *Class 1 (M ± SE)* | *Class 2 (M ± SE)* | *Class 3 (M ± SE)* | *Class 4 (M ± SE)* | *Class 5 (M ± SE)* | *Class 6 (M ± SE)* |
| --- | --- | --- | --- | --- | --- | --- | --- | --- |
| *AUC (ms\*μV)* | **Monophasic** | **PA** | 152.1 ± 25.2 | 177.7 ± 27.9 | 36 ± 22.6 | 88.9 ± 15.5 | 132.1 ± 25.3 | 19.4 ± 16.9 |
|  |  | **AP** | 43.9 ± 18.3 | 199.9 ± 36.5 | 243.8 ± 50.8 | 104 ± 22.7 | 153.4 ± 29.1 | 134 ± 35 |
|  |  | **LM** | 123.8 ± 18.8 | 157.2 ± 31.4 | 41.2 ± 19.1 | 108.6 ± 13.5 | 134.3 ± 25.9 | 20.1 ± 14.4 |
|  | **Biphasic** | **PA** | 83.3 ± 18.2 | 160.8 ± 18.7 | 97.9 ± 31.7 | 135.3 ± 20.6 | 135.1 ± 25 | 33.5 ± 16.9 |
|  |  | **AP** | 93.8 ± 21.9 | 162.4 ± 24.7 | 131 ± 35.3 | 140.3 ± 21.7 | 158.8 ± 20.7 | 53 ± 23.2 |
|  |  | **LM** | 74 ± 15.6 | 166.2 ± 26 | 108.6 ± 18.2 | 163.8 ± 19.8 | 130 ± 23.4 | 52.6 ± 18.4 |
| *Duration (ms)* | **Monophasic** | **PA** | 88.9 ± 10.7 | 114.9 ± 10.3 | 31.4 ± 6.4 | 46.2 ± 6 | 90.2 ± 10.9 | 19.9 ± 4.8 |
|  |  | **AP** | 32.1 ± 6.4 | 102.7 ± 11.1 | 87.6 ± 9.4 | 45 ± 7.5 | 73 ± 10.4 | 53.8 ± 8.5 |
|  |  | **LM** | 84.8 ± 9.5 | 104.5 ± 11.3 | 35.4 ± 7 | 48.5 ± 6.4 | 84.7 ± 11.8 | 22.6 ± 5.3 |
|  | **Biphasic** | **PA** | 52.8 ± 7.5 | 104.6 ± 9.5 | 60.3 ± 7.2 | 61 ± 8.4 | 87.7 ± 11.1 | 27.7 ± 4.5 |
|  |  | **AP** | 51.4 ± 7 | 99.6 ± 9.6 | 67 ± 8.7 | 59 ± 8.3 | 85.5 ± 9.3 | 31.6 ± 5.1 |
| *Onset (ms)* | **Monophasic** | **PA** | 15 ± 3.3 | 55.5 ± 20.3 | 120.2 ± 11.5 | 66.9 ± 6.4 | 71.6 ± 17.2 | 111.6 ± 4.5 |
| | | AP | $65.1 \pm 21.4$ | $173.9 \pm 20$ | $16.4 \pm 5.8$ | $72.1 \pm 12.2$ | $33.4 \pm 14.9$ | $26.1 \pm 7.9$ |
| | | LM | $21.5 \pm 12.6$ | $199.7 \pm 18.7$ | $97.7 \pm 12.5$ | $71.9 \pm 5.8$ | $15.2 \pm 13$ | $90.5 \pm 8.1$ |
| | Biphasic | PA | $38.8 \pm 14.1$ | $94.3 \pm 18.9$ | $11.3 \pm 10.9$ | $57.5 \pm 6$ | $40.6 \pm 18.4$ | $77.6 \pm 12.7$ |
| | | AP | $41.3 \pm 14.1$ | $116 \pm 20.3$ | $25.9 \pm 12.2$ | $37.4 \pm 6$ | $20.8 \pm 9.3$ | $104.8 \pm 9.6$ |
| | | LM | $34.3 \pm 16.7$ | $115.2 \pm 20.2$ | $46.6 \pm 14.8$ | $50.1 \pm 5.6$ | $72.8 \pm 19.8$ | $62.4 \pm 10$ |

#### 3.2.1. Microstate AUC

Changes in the AUC were present only across monophasic conditions, suggesting that microstates were predominant or explained little signal depending on the current direction (for all rmANOVAs main effects and interactions, see **Supplemental Table 4**). Class 1 (*F*1.98,31.73 = 22.40, *p* < .001) had a lower AUC value in AP compared to the other two conditions (PA: *ψ* = -102.11, *p* < .001; LM: *ψ* = -68.76, *p* < .001), and in LM compared to PA (*ψ* = -26.94, *p* < .001, **Figure 6a**). A reversed pattern of modulation was observed both for class 3 (F1.09,17.37 = 20.68, *p* < .001) and class 6 (*F*1.26,20.16 = .26, *p* < .001): in this case, AP currents led to greater AUC than PA (class 3: *ψ* = 189.49, *p* = .025; class 6: *ψ* = 112.97, *p* = .003) and LM ones (class 3: *ψ* = 184.04, *p* < .001; class 6: *ψ* = 117.68, *p* = .002; **Figure 6c** e **f**).

**Figure 6.**
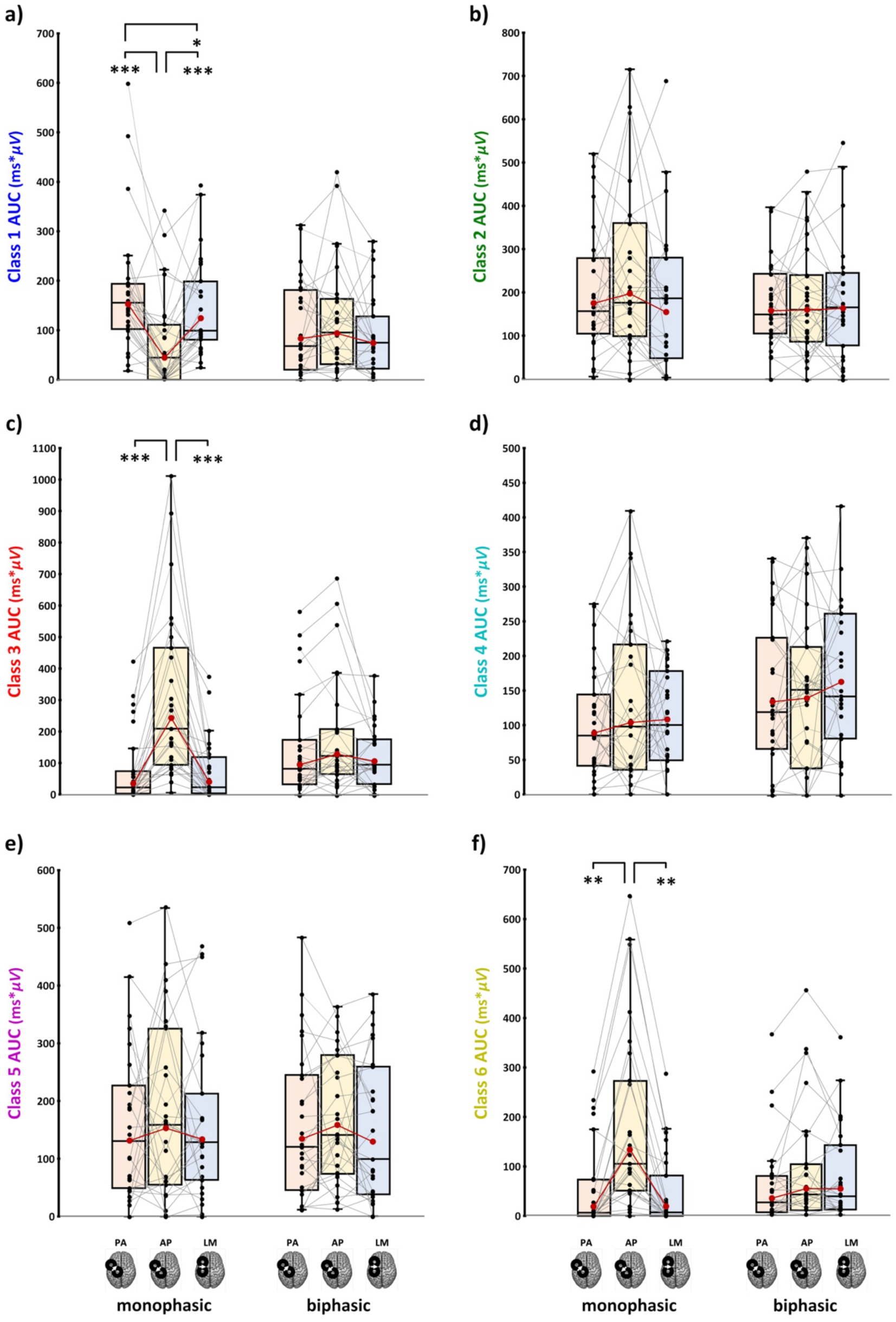
Microstate AUC for all six stimulation conditions for each microstate class. In the box-and-whiskers plots, red dots and lines represent the means of the distributions. The centre line depicts their median values. Black dots and grey lines show individual scores. The box contains the 25th to 75th percentiles of the dataset. Whiskers extend to the largest observation, which falls within the 1.5 times interquartile range from the first/third quartile; significant p-values of corrected post hoc comparisons are reported (* = p <.05; ** = p <.01; *** = p <.001).

For the other three classes, as well as for biphasic conditions, no statistically significant effects were found (all *F*s < 2.56, all *p*s >.092, **Figure 6**).

#### 3.2.2. Microstate Duration

Results on microstate duration were in line with findings on AUC (for all rmANOVAs main effects and interactions, see **Supplemental Table 5**). We found significant ‘Pulse waveform’ X ‘Current direction’ interactions for microstates classes 1, 3, and 6. For class 1 (*F*1.59,41.27 = 16.3, *p* < .001, ƞp^2^ = .39), monophasic AP led to shorter microstates’ duration than the other two monophasic conditions, as well as the homolog biphasic direction. Furthermore, monophasic PA and LM currents evoked longer microstates than the three biphasic directions (for all significant post-hoc comparisons, see **Table 5**, **Figure 7a**). Class 3 (*F*2,52 = 18.3, *p* < .001, ƞp^2^ = .41) showed the opposite pattern found for class 1: namely, monophasic AP evoked microstates with a significantly longer duration than monophasic PA and LM directions whose, in turn, had shorter microstates duration than the three biphasic conditions. In addition, monophasic AP evoked class 3 microstates with a longer duration than biphasic PA and LM directions, suggesting that this current direction evoked the longest class 3 microstates compared to all other conditions except for the homolog biphasic direction (**Figure 7c**). For class 6, monophasic AP led to a longer duration than the other two monophasic conditions, and monophasic PA had a significantly shorter duration than biphasic AP and LM conditions (**Figure 7f**).

**Figure 7.**
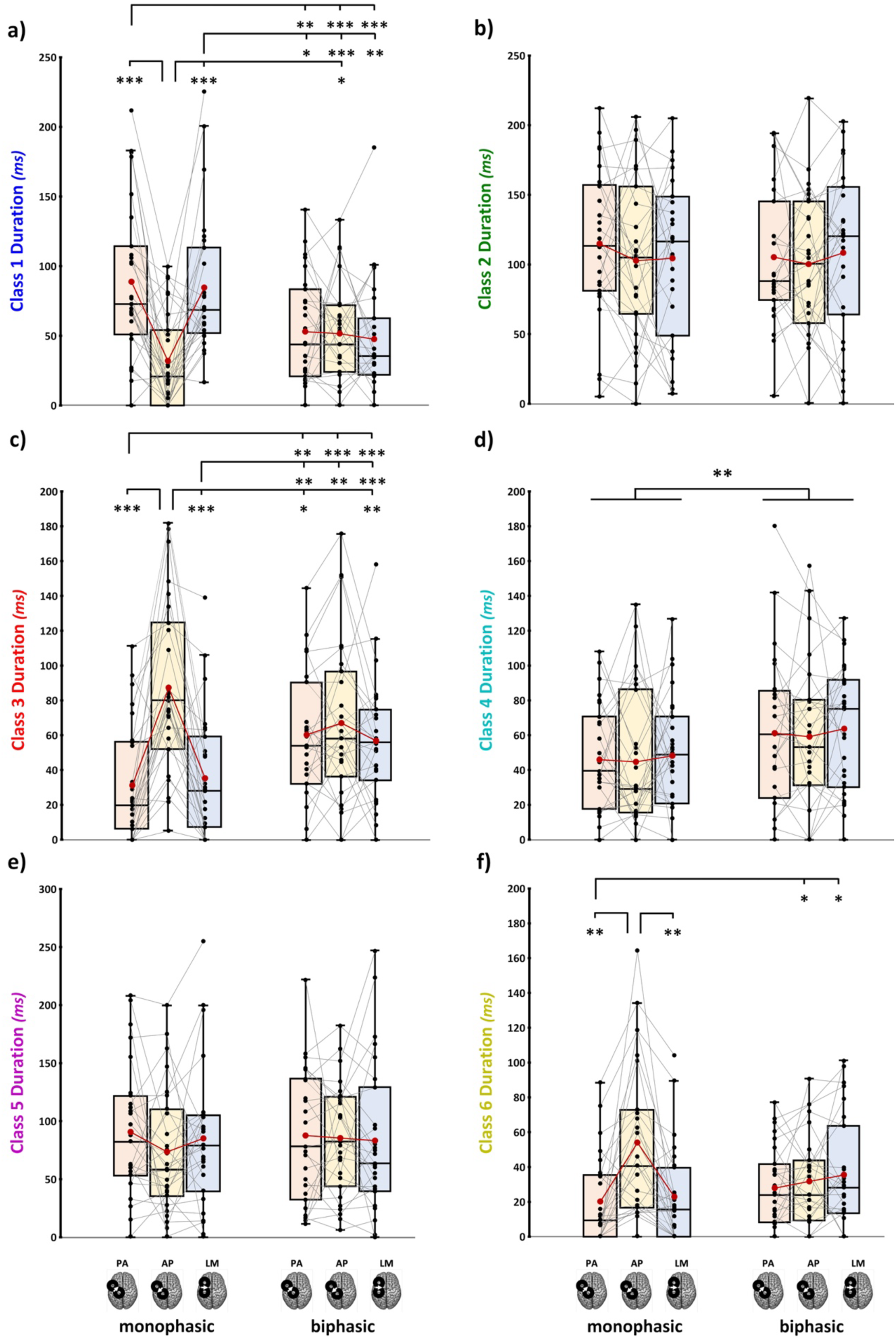
Microstate duration for all six stimulation conditions for each microstate class. In the box-and- whiskers plots, red dots and lines represent the means of the distributions. The centre line depicts their median values. Black dots and grey lines show individual scores. The box contains the 25th to 75th percentiles of the dataset. Whiskers extend to the largest observation, which falls within the 1.5 times interquartile range from the first/third quartile; significant p-values of corrected post hoc comparisons are reported (* = p <.05; ** = p <.01; *** = p <.001).

**Table 5.** Significant post-hoc comparisons (Tukey corrected) for microstate duration.

| *Microstate* | *Condition* | *versus* | *t* | *p<sub>Tukey</sub>* | *Cohen’s d* |
| --- | --- | --- | --- | --- | --- |
| Class 1 | AP<sub>monophasic</sub> | PA<sub>monophasic</sub> | 5.4 | <.001 | 1.06 |
|  |  | LM<sub>monophasic</sub> | 5.1 | <.001 | .99 |
|  |  | AP<sub>biphasic</sub> | -3.2 | .042 | -.61 |
|  | PA<sub>monophasic</sub> | PA<sub>biphasic</sub> | 4 | .005 | .78 |
|  |  | AP<sub>biphasic</sub> | 4.9 | <.001 | .94 |
|  |  | LM<sub>biphasic</sub> | 4.8 | <.001 | .91 |
|  | LM<sub>monophasic</sub> | PA<sub>biphasic</sub> | 3.2 | .042 | .61 |
|  |  | AP<sub>biphasic</sub> | 4.8 | <.001 | .92 |
|  |  | LM<sub>biphasic</sub> | 4 | .005 | .78 |
| **Class 3** | **AP<sub>monophasic</sub>** | **PA<sub>monophasic</sub>** | -6.6 | <.001 | -1.26 |
|  |  | **AP<sub>monophasic</sub>** | -5.5 | <.001 | -1.06 |
|  |  | **PA<sub>biphasic</sub>** | 3.8 | .01 | .72 |
|  |  | **LM<sub>biphasic</sub>** | 3.4 | .002 | .66 |
|  | **PA<sub>monophasic</sub>** | **PA<sub>biphasic</sub>** | -4.7 | .001 | -.9 |
|  |  | **AP<sub>biphasic</sub>** | -4.8 | <.001 | -.92 |
|  |  | **LM<sub>biphasic</sub>** | -4.9 | <.001 | -.94 |
|  | **LM<sub>monophasic</sub>** | **PA<sub>biphasic</sub>** | -3.9 | .006 | -.77 |
|  |  | **AP<sub>biphasic</sub>** | 4.1 | .004 | -.8 |
|  |  | **LM<sub>biphasic</sub>** | -4.3 | <.001 | -.82 |
| **Class 6** | **AP<sub>monophasic</sub>** | **AP<sub>monophasic</sub>** | -4.2 | .004 | .81 |
|  |  | **LM<sub>monophasic</sub>** | 4 | .005 | .77 |
|  | **PA<sub>monophasic</sub>** | **AP<sub>biphasic</sub>** | -3.2 | .04 | -.61 |
|  |  | **LM<sub>biphasic</sub>** | -3.4 | .02 | -.66 |

Notably, for class 4, a significant main effect of ‘Pulse waveform’ was found: regardless of the current direction exploited, microstates found with monophasic waveforms had shorter durations than the ones found with biphasic pulses (*F*1,26 = 11.7, *p* = .002, ƞp^2^ = .31, **Figure 7d**).

Finally, for classes 2 and 5, no significant differences were found (all *F*s < 1.8, all *p*s >.176, **Figure 7**).

#### 3.2.3. Microstate order of appearance

Results on microstate onset showed a significant main effect of ‘Microstate class’ (*χ²*5 = 82.91, *p <* 0.001) as well as a significant interaction between ‘Microstate class’ and ‘Stimulation condition’ (*χ²*25 = 141.54, *p* < .001). Specifically, the interaction effect showed changes in microstates’ order of appearance across conditions, with four significant inversions of order between microstate classes across conditions. These swaps in microstates onset should not be interpreted as modulations of the timing at which a given neural process occurs, e.g., it is improbable that the same neural process usually observed at the late latency suddenly happens at the very beginning, or vice-versa. Rather, it is more likely that a new neural process characterized by the same topography occurs earlier. Importantly, these swaps indicate that the transitions between microstates do not follow the same order across conditions, suggesting distinct effective connectivity patterns.

First, two classes swapped between biphasic and monophasic AP conditions. While in biphasic AP the onset of classes 4 and 5 significantly preceded the onset of class 6 (class 4 vs. class 6: *p =* .023; class 5 vs. class 6: *p=* .006), the order was reversed in monophasic AP, such that classes 4 and 5 followed class 6 (class 4 vs. class 6: *p =* .003; class 5 vs. class 6: *p=* .013). Then, classes 1, 3 and 6 reversed their relative order across monophasic conditions: while class 3 and 6 significantly preceded class 1 in monophasic AP (class 1 vs. class 3: *p =* .004; class 1 vs. class 6: *p <* .001), class 1 appeared before both class 3 (LM: *p <* .001; PA: *p* < .001) and class 6 (LM: *p =* .003; PA: *p <* .001; **Figure 8**) in monophasic LM and PA.

**Figure 8.**
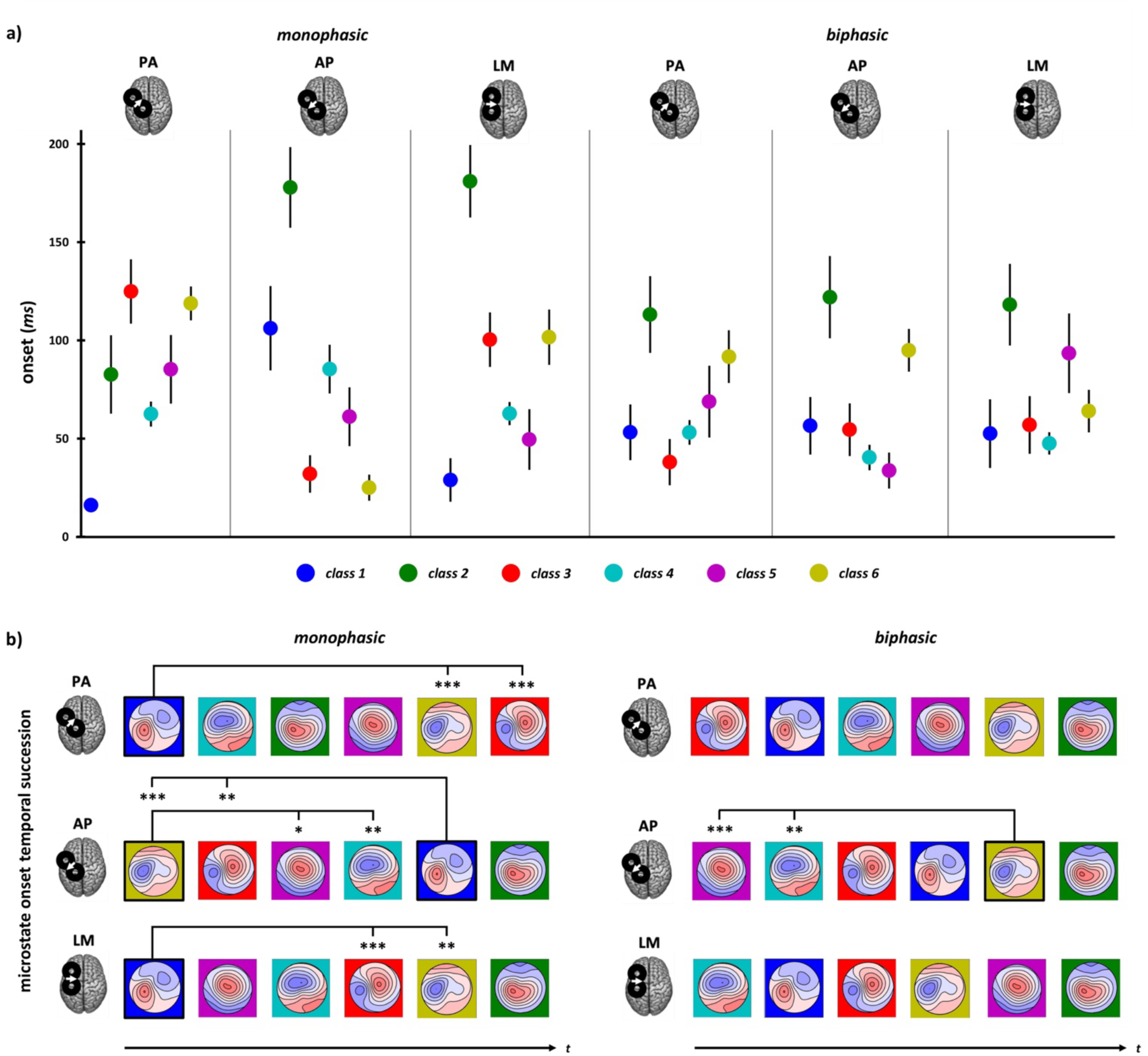
(a) Mean microstate onset of the six different classes in the six experimental blocks. Error bars represent standard error. **(b)** Temporal succession of microstate onset classes in the different stimulation conditions. Significant swaps of class appearance across conditions are reported (* = p <.05; ** = p <.01; *** = p <.001).

All significant differences in microstates’ onset, including contrasts not involving swaps between classes across conditions, are reported in **Table 6**.

**Table 6.** Significant post-hoc comparisons between microstate classes’ onset within each stimulation condition. Contrasts highlighted in bold indicate those involved in a swap in the order of appearance across stimulation conditions. Reported p-values are corrected for multiple comparisons using the Benjamini-Hochberg method, considering 90 contrasts of interest (15 possible contrasts between classes across 6 stimulation conditions).

| *Stimulation Condition* | *Microstate classes comparison* | *Estimate* | *p* |
| --- | --- | --- | --- |
| **Monophasic PA** | 1 vs. 2 | -67.3 | <.001 |
|  | **1 vs. 3** | **-110.7** | **<.001** |
|  | 1 vs. 4 | -48.8 | <.001 |
|  | 1 vs. 5 | -71.2 | <.001 |
|  | **1 vs. 6** | **-103.1** | **<.001** |
| **Monophasic AP** | 1 vs. 2 | -74.0 | .043 |
|  | **1 vs. 3** | **70.3** | **.004** |
|  | **1 vs. 6** | **77.7** | **<.001** |
|  | 2 vs. 3 | 144.3 | <.001 |
|  | 2 vs. 4 | 92.5 | .008 |
|  | 2 vs. 5 | 116.1 | <.001 |
|  | 2 vs. 6 | 151.8 | <.001 |
|  | 3 vs. 4 | -51.8 | .011 |
|  | **4 vs. 6** | **59.2** | **.003** |
|  | **5 vs. 6** | **35.6** | **.013** |
| **Monophasic LM** | 1 vs. 2 | -159.3 | <.001 |
|  | **1 vs. 3** | **-78.1** | **<.001** |
|  | 1 vs. 4 | -41.1 | .006 |
|  | 1 vs. 5 | -24.4 | .045 |
|  | **1 vs. 6** | **-78.9** | **.003** |
|  | 2 vs. 4 | 118.1 | <.001 |
|  | 2 vs. 5 | 134.9 | <.001 |
|  | 3 vs. 5 | 53.7 | .03 |
| **Biphasic PA** | 1 vs. 2 | -65.8 | .009 |
|  | 2 vs. 3 | 78.3 | <.001 |
|  | 2 vs. 4 | 60.2 | .016 |
|  | 3 vs. 6 | -54.9 | .014 |
| **Biphasic AP** | 1 vs. 2 | -70.7 | .006 |
|  | 2 vs. 3 | 66.7 | .009 |
|  | 2 vs. 4 | 80.9 | <.001 |
|  | 2 vs. 5 | 90.0 | <.001 |
|  | **4 vs. 6** | **-55.5** | **.023** |
|  | **5 vs. 6** | **-64.6** | **.006** |
| **Biphasic LM** | 1 vs. 2 | -63.2 | .023 |
|  | 2 vs. 3 | 60.9 | .026 |
|  | 2 vs. 4 | 69.9 | .007 |
|  | 4 vs. 5 | -45.4 | .043 |

## 4. DISCUSSION

Our results show evidence that manipulating TMS parameters evokes activation of distinct cortical pathways within the stimulated network, thereby generating variability in the response to stimulation. The same M1-TEP components and microstate maps are observed across our stimulation conditions but with different spatiotemporal patterns, revealing a modulation of M1 signal propagation that is more pronounced for monophasic pulses than biphasic ones.

### 4.1. TMS pulse waveform and current direction affect M1 effective connectivity

Considering the general pattern found for amplitude, latency, and topographical profiles of TEP data, the first striking evidence is that monophasic waveform affects the amplitude of all the analyzed M1- TEP components compared to biphasic waveforms. At the same time, biphasic stimulation leads to changes among current directions only for a couple of components (i.e., P30 and P60). This pattern is not surprising considering the physical characteristics of electric current flow in the brain elicited by TMS: indeed, for monophasic pulses, there is only one phase resulting in neuronal stimulation, while biphasic pulses have two physiologically effective phases, with the second one being more efficacious in neuronal excitation than the first one (e.g., Corthout et al., 2001; Groppa et al., 2012; Maccabee et al., 1998). Hence, monophasic pulses induce higher direction specificity for the stimulated neuronal population (Sommer et al., 2018), likely allowing the activation of more selective circuits than biphasic ones.

Microstate results are particularly informative in this regard by depicting cortical activity spatiotemporal distribution. Similar to TEP amplitude and latency patterns, the differences observed between current directions in microstate succession and their parameters are more evident for monophasic conditions than biphasic ones, suggesting more pronounced changes in effective connectivity across monophasic stimulation conditions. In detail, analysis of microstate duration and AUC showed that monophasic waveforms lead to greater modulations compared to biphasic ones, with AP direction significantly differing from PA and LM ones for microstate classes 1, 3, and 6. The analysis on microstate onset enriched this evidence, showing that inversion of the order of classes’ appearance is prominent only within monophasic conditions and for microstates presenting AUC and duration modulations. Namely, class 1 appearance occurs significantly later during AP stimulation than classes 3 and 6. At the same time, the reverse pattern is seen with PA and LM stimulations (i.e., class 1 appearance occurred earlier than classes 3 and 6). Furthermore, concerning homologous current directions, only monophasic *versus* biphasic AP direction shows statistically significant changes in microstate’s order, with class 6 preceding the occurrence of classes 4 and 5 during monophasic AP stimulation but succeeding them during biphasic AP, pointing out that AP direction is the one activating the most dissimilar network profile also between pulse waveforms. As stated previously, these swaps in microstate order likely indicates the appearance of new neural processes with similar topography rather than timing changes in existing processes. Therefore, these changes in the sequence of transitions from one microstate to another likely reflect different patterns of effective connectivity. Altogether, this evidence suggests that utilizing biphasic waveforms engages a widespread activity from M1, reflecting the contribution of cortical networks that could be selectively activated only when a specific current direction is employed, as with monophasic waveforms.

Another result stemming from the overall TEP patterns is the more pronounced impact of our experimental manipulations on TEP amplitudes rather than their latencies, suggesting that the propagation speed across neuronal networks is largely unaffected by TMS parameters (but see effects on N15 and on P15 previously reported, e.g., Guidali et al., 2023; Bonato et al., 2006).

Finally, TMS parameters appear to have a greater influence on earlier TEP components, corroborating previous studies showing technical parameter modulations specifically for early cortical responses (Bonato et al., 2006; Casula et al., 2018). In detail, in our sample, many participants show an inverted polarity or a lack of N15, P30, and N45 components, while later peaks only exhibit an amplitude modulation. This evidence is also highlighted by the overall pattern of microstate class succession, where a substantial modulation among conditions (and, in detail, among monophasic ones) is observed within the first 50 ms after TMS.

Altogether, these results suggest the activation of distinct cortical circuits and/or differential engagement of similar cortical circuits when different TMS parameters are used.

### 4.2. Monophasic anterior-posterior currents maximize the recruitment of distinct motor circuitries in the first 50 ms

As presented before, the monophasic AP condition shows greater differences for TEP peaks and microstate parameters than all the other experimental conditions. Notably, classes 6 and 3 topographies, which represent the first microstates found after TMS in monophasic AP stimulation, are characterised by prominent negativity under the stimulation site reflecting the N15 component, thought to disclose the activation of sensorimotor and premotor areas ipsilateral to stimulation (Farzan & Bortoletto, 2022). Also, class 3 shows a positivity over sensorimotor electrodes contralateral to stimulation, resembling an early component linked to the callosal inhibition of the contralateral M1 (M1-P15; Bortoletto et al. 2021; Zazio et al. 2022) having a higher amplitude when elicited with monophasic AP pulses (Guidali et al., 2023). Hence, if N15 amplitude and latency tell us that monophasic AP is the condition that permits recruiting the neuronal populations involved more effectively and rapidly, microstates analysis shows that this condition also activates different cortical circuits than other monophasic pulses, as reflected by different EEG topographies within the first 50 ms after TMS. Furthermore, it allows us to hypothesise that AP monophasic pulse preferentially engages ipsilateral and cortico-cortical motor populations. This pattern is further confirmed by class 1 modulations, i.e., the microstate presenting the lowest duration and AUC during monophasic AP stimulation and characterising the first responses to monophasic LM and PA. Indeed, its topography, with positivity over the stimulation site and negativity over contralateral frontocentral electrodes, closely reassembles N45 topography, which, in turn, is the only TEP component that is significantly reduced after monophasic AP stimulation.

Previous literature suggested changes in the spatial selectivity induced by the two PA and AP current directions when stimulating M1 (e.g., Aberra et al., 2020; Siebner et al., 2022; Spampinato, 2020), and our results further corroborate this evidence. AP current direction is thought to lead to an anterior shift of the maximal TMS-induced electric field amplitude, thereby stimulating neurons located more anteriorly in the precentral gyrus compared to PA (and, to a lesser extent, LM) current direction (Aberra et al., 2020). Rostral neurons of the precentral gyrus are more interconnected with higher-order motor areas (e.g., supplementary motor area) than caudal ones (Spampinato, 2020). Thus, the AP current direction can stimulate higher-order motor populations more efficiently than PA and LM directions, leading to a different spread of M1 activation. Furthermore, besides better activating the rostral part of the precentral gyrus, previous TMS studies involving repetitive and paired-pulse protocols pointed out that AP currents more efficiently stimulate M1 superficial layers (i.e., L2/3), while PA and LM currents reach deeper layers (i.e., L5) more easily (e.g., Casarotto et al., 2023; Koch et al., 2013; Sommer et al., 2013). Interestingly, studies on animal models highlighted that M1 interneurons responsible for cortico-cortical communication and network-wide activations are more numerous in superficial layers of the motor cortex rather than in deeper ones, where instead are predominant neuronal population directly activating the pyramidal tract (e.g., Harris & Shepherd, 2015; Mao et al., 2011; Weiler et al., 2008). Hence, AP stimulation could be more effective in directly activating cortico-cortical neurons and, in turn, cortical regions structurally interconnected with M1, as supported by our microstates and TEP results.

### 4.3. Modulation of M1-TEP components associated with inhibitory and excitatory circuits

Modulation patterns found for TEP components associated with inhibitory and excitatory circuits corroborate further the evidence that TMS parameters modulate M1 effective connectivity. Considering negative components, different pharmacologic studies suggest that N45 and N100 are related to gamma-aminobutyric acid (GABA) circuitries (Darmani & Ziemann, 2019; Premoli, Castellanos, et al., 2014). Notably, the N45 is linked to inhibitory processes mediated by GABA-A receptors (Belardinelli et al., 2021; Cash et al., 2017; Darmani et al., 2016), while the N100 is associated with GABA-B ones (Premoli et al., 2018; Premoli, Rivolta, et al., 2014; Rogasch et al., 2013). Our results show that monophasic AP elicits the lowest N45 amplitude but the highest N100 amplitude compared to PA and LM. This could imply that when monophasic waveforms are employed, AP and PA/LM directions can activate cortical circuits to different extents, permitting the selective modulation of GABA-A-mediated (the latter) or GABA-B-mediated (the former) pathways. Further investigating this possibility would be extremely useful for clinical purposes due to GABAergic dysfunctions associated with different stages or specific symptoms of numerous clinical the disorders (e.g., Alzheimer’s disease, Major Depressive disorder, schizophrenia; Heaney & Kinney, 2016; Luscher et al., 2011).

The differences in positive TEP components observed across conditions are less straightforward to explain. The P30 is presumed to reflect excitatory activity with involvement from multiple cortical sources (Farzan & Bortoletto, 2022), and it has been associated with MEP amplitude (e.g., Ahn & Fröhlich, 2021; Mäki & Ilmoniemi, 2010). Interestingly, previous studies showed that medial-lateral direction is one of the least effective directions for evoking MEPs (e.g., Mills et al., 1992; Souza et al., 2018). Thus, when the lateral-medial current direction is employed with a biphasic pulse, it may impact M1 cortical reactivity, leading to the pattern observed in our P30 data.

Similar results were found for the P60, where LM is still the current direction evoking the smaller components overall. Previous TMS-EEG literature linked P60 amplitude to the cortical processing of the MEP sensorimotor reafference (Mäki & Ilmoniemi, 2010; Petrichella et al., 2017). Crucially, in contrast with TEP findings, MEP amplitude registered in the current dataset showed no differences among stimulation conditions (see Guidali et al., 2023 and **Supplemental Figure 2**). Hence, P60 patterns could rather reflect an intermingled effect of muscular reafferent elaboration (e.g., no differences between PA and AP directions) and cortico-cortical activity (e.g., LM lower efficiency).

Finally, considering the contribution of cortical processes not strictly related to the direct stimulation of M1, the latest component we investigated, the P180, is the only one showing a significant contribution of TMS intensity for both pulse waveforms. This result is in line with the current literature highlighting the contribution of TMS-related auditory and somatosensory inputs on this component, whose activity is located mainly over temporoparietal regions (Biabani et al., 2019; Conde et al., 2019). Indeed, it is reasonable to think that the higher rMT found for monophasic AP stimulation (see **Supplemental Figure 1**) – and, consequently, the higher stimulation intensity, had caused in our participants stronger somatosensory sensations on the scalp and a louder TMS noise (i.e., the ‘click’ sound – which our white noise could not entirely mask), leading to greater somatosensory and auditory evoked potentials.

It must be noted that the relation between later TEP components and TMS-related artifacts has also been previously demonstrated for the N100 (Biabani et al., 2019). Hence, even if our data did not show any role of stimulation intensity on this negative component, we cannot entirely exclude the contamination of N100 with auditory and somatosensory processing.

### 4.4. Future directions and limitations

The present study has a few limitations that must be taken into account. First, microstate variables showed a high inter-individual variability (see **Figures 6** and **7**); therefore, our conclusions on brain network modulations must be interpreted cautiously (e.g., Kleinert et al., 2024). Then, other TMS parameters than those we considered, such as TMS pulse width, can impact brain responses to stimulation (Casula et al., 2018) and should be included in future studies in combination with the parameters described here. Finally, stimulation intensity was based on the participant’s rMT, which, on the one hand, helped us control that all conditions elicited the same corticospinal activity but, on the other hand, did not ensure comparable cortical reactivity across conditions.

Future TMS-EEG studies should then carefully consider TMS parameters in order to use TEPs as biomarkers in a reliable way. Moreover, expanding this knowledge could help optimize modulatory TMS protocols acting or influencing motor system cortico-cortical connectivity, permitting the better targeting of specific neural pathways/circuits, especially when these protocols are used in clinical settings (Guidali et al., 2021; Hernandez-Pavon, San Agustín, et al., 2023). Again, given the promising results obtained at the sensor level, source-level connectivity analysis could deepen the present results, going beyond the information conveyed by microstate analysis. Last but not least, this investigation could be extended to other cortical areas (e.g., dorsolateral prefrontal cortex, inferior parietal lobule) known to play a crucial role in neuropsychiatric disorders (Cao et al., 2021; Farzan, 2024).

## 5. CONCLUSION

To conclude, our study highlights the pivotal role of TMS parameters over TEP modulation and their variability, suggesting that different cortical circuits and networks are recruited according to the current direction and pulse waveform chosen. Overall, biphasic stimulation allows the change of coil orientation without highly affecting the evoked response, while monophasic one helps achieve a higher stimulation selectivity, as supported by other TMS studies (e.g., Fong et al., 2021; Hamada et al., 2014; Pieramico et al., 2023; Sale et al., 2016). Critically, AP current direction led to the most dissimilar spatiotemporal patterns compared to the other conditions exploited. Thus, depending on the aim of the study, the most suitable TMS parameters must be carefully chosen, given that they are an important source of variability when TMS-EEG is used. Furthermore, considering microstate analysis applied to TMS-EEG data (Ding et al., 2024; Sulcova et al., 2022), our results suggest that they could provide valuable information at the sensor level, likely complementary to the one obtained by solely looking at amplitude and latency of evoked responses, helping infer the activation of distinct cortical circuits.

## CRedIT AUTHOR CONTRIBUTION

**Delia Lucarelli:** conceptualization, methodology, investigation, formal analysis, software, visualization, writing – original draft

**Giacomo Guidali:** conceptualization, methodology, investigation, formal analysis, data curation, visualization, writing – original draft

**Dominika Sulcova:** methodology, writing – review and editing

**Agnese Zazio:** formal analysis, visualization, writing – review and editing

**Natale Salvatore Bonfiglio**: formal analysis, writing – review and editing **Antonietta Stango:** software, writing – review and editing

**Guido Barchiesi:** formal analysis, writing – review and editing

**Marta Bortoletto:** conceptualization, methodology, supervision, validation, resources, funding acquisition, writing – original draft

## REFERENCES

1. Aberra, A. S., Wang, B., Grill, W. M., & Peterchev, A. V. (2020). Simulation of transcranial magnetic stimulation in head model with morphologically-realistic cortical neurons. Brain Stimulation, 13(1), 175–189. 10.1016/J.BRS.2019.10.002

2. Ahn, S., & Fröhlich, F. (2021). Pinging the brain with transcranial magnetic stimulation reveals cortical reactivity in time and space. Brain Stimulation, 14(2), 304–315. 10.1016/j.brs.2021.01.018

3. Awiszus, F. (2003). TMS and threshold hunting. Supplements to Clinical Neurophysiology, 56(C), 13–23. 10.1016/S1567-424X(09)70205-3

4. Bagattini, C., Mutanen, T. P., Fracassi, C., Manenti, R., Cotelli, M., Ilmoniemi, R. J., Miniussi, C., & Bortoletto, M. (2019). Predicting Alzheimer’s disease severity by means of TMS–EEG coregistration. Neurobiology of Aging, 80, 38–45. 10.1016/j.neurobiolaging.2019.04.008

5. Bates, D., Mächler, M., Bolker, B. M., & Walker, S. C. (2015). Fitting linear mixed-effects models using lme4. Journal of Statistical Software, 67(1). 10.18637/jss.v067.i01

6. Beck, M. M., Heyl, M., Mejer, L., Vinding, M. C., Christiansen, L., Tomasevic, L., & Siebner, H. R. (2024). Methodological Choices Matter: A Systematic Comparison of TMS-EEG Studies Targeting the Primary Motor Cortex. Human Brain Mapping, 45(15). 10.1002/hbm.70048

7. Belardinelli, P., König, F., Liang, C., Premoli, I., Desideri, D., Müller-Dahlhaus, F., Gordon, P. C., Zipser, C., Zrenner, C., & Ziemann, U. (2021). TMS-EEG signatures of glutamatergic neurotransmission in human cortex. Scientific Reports, 11(1), 1–14. 10.1038/s41598-021-87533-z

8. Biabani, M., Fornito, A., Mutanen, T. P., Morrow, J., & Rogasch, N. C. (2019). Characterizing and minimizing the contribution of sensory inputs to TMS-evoked potentials. Brain Stimulation, 12(6), 1537–1552. 10.1016/j.brs.2019.07.009

9. Bonato, C., Miniussi, C., & Rossini, P. M. (2006). Transcranial magnetic stimulation and cortical evoked potentials: A TMS/EEG co-registration study. Clinical Neurophysiology, 117(8), 1699–1707. 10.1016/j.clinph.2006.05.006

10. Bortoletto, M., Bonzano, L., Zazio, A., Ferrari, C., Pedullà, L., Gasparotti, R., Miniussi, C., & Bove, M. (2021). Asymmetric transcallosal conduction delay leads to finer bimanual coordination. Brain Stimulation, 14. 10.1016/j.brs.2021.02.002

11. Bortoletto, M., Veniero, D., Thut, G., & Miniussi, C. (2015). The contribution of TMS-EEG coregistration in the exploration of the human cortical connectome. Neuroscience and Biobehavioral Reviews, 49, 114–124. 10.1016/j.neubiorev.2014.12.014

12. Cao, K. X., Ma, M. L., Wang, C. Z., Iqbal, J., Si, J. J., Xue, Y. X., & Yang, J. L. (2021). TMS-EEG: An emerging tool to study the neurophysiologic biomarkers of psychiatric disorders. Neuropharmacology, 197, 108574. 10.1016/J.NEUROPHARM.2021.108574

13. Casarotto, A., Dolfini, E., Cardellicchio, P., Fadiga, L., D’Ausilio, A., & Koch, G. (2023). Mechanisms of Hebbian-like plasticity in the ventral premotor – primary motor network. The Journal of Physiology, 601(1), 211–226. 10.1113/JP283560

14. Casarotto, S., Turco, F., Comanducci, A., Perretti, A., Marotta, G., Pezzoli, G., Rosanova, M., & Isaias, I. U. (2019). Excitability of the supplementary motor area in Parkinson’s disease depends on subcortical damage. Brain Stimulation, 12(1), 152–160. 10.1016/j.brs.2018.10.011

15. Cash, R. F. H., Noda, Y., Zomorrodi, R., Radhu, N., Farzan, F., Rajji, T. K., Fitzgerald, P. B., Chen, R., Daskalakis, Z. J., & Blumberger, D. M. (2017). Characterization of Glutamatergic and GABA A-Mediated Neurotransmission in Motor and Dorsolateral Prefrontal Cortex Using Paired-Pulse TMS-EEG. Neuropsychopharmacology, 42(2), 502–511. 10.1038/npp.2016.133

16. Casula, E. P., Borghi, I., Maiella, M., Pellicciari, M. C., Bonnì, S., Mencarelli, L., Assogna, M., D’Acunto, A., Di Lorenzo, F., Spampinato, D. A., Santarnecchi, E., Martorana, A., & Koch, G. (2023). Regional Precuneus Cortical Hyperexcitability in Alzheimer’s Disease Patients. Annals of Neurology, 93(2), 371–383. 10.1002/ana.26514

17. Casula, E. P., Rocchi, L., Hannah, R., & Rothwell, J. C. (2018). Effects of pulse width, waveform and current direction in the cortex: A combined cTMS-EEG study. Brain Stimulation, 11(5), 1063–1070. 10.1016/j.brs.2018.04.015

18. Cirillo, J., & Byblow, W. D. (2016). Threshold tracking primary motor cortex inhibition: the influence of current direction. European Journal of Neuroscience, 44(8), 2614–2621. 10.1111/ejn.13369

19. Conde, V., Tomasevic, L., Akopian, I., Stanek, K., Saturnino, G. B., Thielscher, A., Bergmann, T. O., & Siebner, H. R. (2019). The non-transcranial TMS-evoked potential is an inherent source of ambiguity in TMS-EEG studies. NeuroImage, 185(September 2018), 300–312. 10.1016/j.neuroimage.2018.10.052

20. Corp, D. T., Bereznicki, H. G. K., Clark, G. M., Youssef, G. J., Fried, P. J., Jannati, A., Davies, C. B., Gomes-Osman, J., Kirkovski, M., Albein-Urios, N., Fitzgerald, P. B., Koch, G., Di Lazzaro, V., Pascual-Leone, A., & Enticott, P. G. (2021). Large-scale analysis of interindividual variability in single and paired-pulse TMS data. Clinical Neurophysiology, xxxx. 10.1016/j.clinph.2021.06.014

21. Corthout, E., Barker, A. T., & Cowey, A. (2001). Transcranial magnetic stimulation: Which part of the current waveform causes the stimulation? Experimental Brain Research, 141(1), 128–132. 10.1007/s002210100860

22. Darmani, G., & Ziemann, U. (2019). Pharmacophysiology of TMS-evoked EEG potentials: A mini-review. Brain Stimulation, 12(3), 829–831. 10.1016/j.brs.2019.02.021

23. Darmani, G., Zipser, C. M., Böhmer, G. M., Deschet, K., Müller-Dahlhaus, F., Belardinelli, P., Schwab, M., & Ziemann, U. (2016). Effects of the selective α5-GABAAR antagonist S44819 on excitability in the human brain: A TMS– EMG and TMS–EEG phase I study. Journal of Neuroscience, 36(49), 12312–12320. 10.1523/JNEUROSCI.1689-16.2016

24. Davila-Pérez, P., Jannati, A., Fried, P. J., Cudeiro Mazaira, J., & Pascual-Leone, A. (2018). The Effects of Waveform and Current Direction on the Efficacy and Test–Retest Reliability of Transcranial Magnetic Stimulation. Neuroscience, 393, 97–109. 10.1016/j.neuroscience.2018.09.044

25. Delorme, A., & Makeig, S. (2004). EEGLAB: an open source toolbox for analysis of single-trial EEG dynamics including independent component analysis. Journal of Neuroscience Methods, 134(1), 9–21. 10.1016/J.JNEUMETH.2003.10.009

26. Delvendahl, I., Lindemann, H., Jung, N. H., Pechmann, A., Siebner, H. R., & Mall, V. (2014). Influence of waveform and current direction on Short-interval intracortical facilitation: A Paired-pulse TMS study. Brain Stimulation, 7(1), 49–58. 10.1016/j.brs.2013.08.002

27. Di Lazzaro, V., Oliviero, A., Saturno, E., Pilato, F., Insola, A., Mazzone, P., Profice, P., Tonali, P., & Rothwell, J. C. (2001). The effect on corticospinal volleys of reversing the direction of current induced in the motor cortex by transcranial magnetic stimulation. Experimental Brain Research, 138(2), 268–273. 10.1007/s002210100722

28. Di Lazzaro, V., Rothwell, J., & Capogna, M. (2018). Noninvasive Stimulation of the Human Brain: Activation of Multiple Cortical Circuits. Neuroscientist, 24(3), 246–260. 10.1177/1073858417717660

29. Ding, Z., Wang, Y., Niu, Z., Ouyang, G., & Li, X. (2024). The effect of EEG microstate on the characteristics of TMS-EEG. Computers in Biology and Medicine, 173(January), 108332. 10.1016/j.compbiomed.2024.108332

31. D’Ostilio, K., Goetz, S. M., Hannah, R., Ciocca, M., Chieffo, R., Chen, J. C. A., Peterchev, A. V., & Rothwell, J. C. (2016). Effect of coil orientation on strength-duration time constant and I-wave activation with controllable pulse parameter transcranial magnetic stimulation. Clinical Neurophysiology, 127(1), 675–683. 10.1016/j.clinph.2015.05.017

32. Farzan, F. (2024). Transcranial Magnetic Stimulation–Electroencephalography for Biomarker Discovery in Psychiatry. Biological Psychiatry, 95(6), 564–580. 10.1016/j.biopsych.2023.12.018

33. Farzan, F., Barr, M. S., Hoppenbrouwers, S. S., Fitzgerald, P. B., Chen, R., Pascual-Leone, A., & Daskalakis, Z. J. (2013). The EEG correlates of the TMS-induced EMG silent period in humans. NeuroImage, 83, 120–134. 10.1016/j.neuroimage.2013.06.059

34. Farzan, F., & Bortoletto, M. (2022). Identification and verification of a “true” TMS evoked potential in TMS-EEG. Journal of Neuroscience Methods, 378(October 2021), 109651. 10.1016/j.jneumeth.2022.109651

35. Federico, P., & Perez, M. A. (2017). Distinct Corticocortical Contributions to Human Precision and Power Grip. Cerebral Cortex, 27(11), 5070–5082. 10.1093/cercor/bhw291

36. Fong, P. Y., Spampinato, D., Rocchi, L., Hannah, R., Teng, Y., Di Santo, A., Shoura, M., Bhatia, K., & Rothwell, J. C. (2021). Two forms of short-interval intracortical inhibition in human motor cortex. Brain Stimulation, 14(5), 1340–1352. 10.1016/j.brs.2021.08.022

37. Gefferie, S. R., Jiménez-Jiménez, D., Visser, G. H., Helling, R. M., Sander, J. W., Balestrini, S., & Thijs, R. D. (2023). Transcranial magnetic stimulation-evoked electroencephalography responses as biomarkers for epilepsy: A review of study design and outcomes. Human Brain Mapping, 44(8), 3446–3460. 10.1002/hbm.26260

38. Gordon, P. C., Jovellar, D. B., Song, Y. F., Zrenner, C., Belardinelli, P., Siebner, H. R., & Ziemann, U. (2021). Recording brain responses to TMS of primary motor cortex by EEG – utility of an optimized sham procedure. NeuroImage, 245(October), 118708. 10.1016/j.neuroimage.2021.118708

39. Groppa, S., Schlaak, B. H., Münchau, A., Werner-Petroll, N., Dünnweber, J., Bäumer, T., van Nuenen, B. F. L., & Siebner, H. R. (2012). The human dorsal premotor cortex facilitates the excitability of ipsilateral primary motor cortex via a short latency cortico-cortical route. Human Brain Mapping, 33(2), 419–430. 10.1002/hbm.21221

40. Guidali, G., Roncoroni, C., & Bolognini, N. (2021). Paired associative stimulations: Novel tools for interacting with sensory and motor cortical plasticity. Behavioural Brain Research, 414(September), 113484. 10.1016/j.bbr.2021.113484

41. Guidali, G., Zazio, A., Lucarelli, D., Marcantoni, E., Stango, A., Barchiesi, G., & Bortoletto, M. (2023). Effects of transcranial magnetic stimulation (TMS) current direction and pulse waveform on cortico-cortical connectivity: A registered report TMS-EEG study. European Journal of Neuroscience, 58(8), 3785–3809. 10.1111/ejn.16127

42. Habermann, M., Weusmann, D., Stein, M., & Koenig, T. (2018). A student’s guide to randomization statistics for multichannel event-related potentials using Ragu. Frontiers in Neuroscience, 12(JUN). 10.3389/fnins.2018.00355

43. Hamada, M., Galea, J. M., Di Lazzaro, V., Mazzone, P., Ziemann, U., & Rothwell, J. C. (2014). Two Distinct Interneuron Circuits in Human Motor Cortex Are Linked to Different Subsets of Physiological and Behavioral Plasticity. Journal of Neuroscience, 34(38), 12837–12849. 10.1523/JNEUROSCI.1960-14.2014

44. Harris, K. D., & Shepherd, G. M. G. (2015). The neocortical circuit: Themes and variations. Nature Neuroscience, 18(2), 170–181. 10.1038/nn.3917

45. Heaney, C. F., & Kinney, J. W. (2016). Role of GABAB receptors in learning and memory and neurological disorders. In Neuroscience and Biobehavioral Reviews (Vol. 63, pp. 1–28). Elsevier Ltd. 10.1016/j.neubiorev.2016.01.007

46. Hernandez-Pavon, J. C., San Agustín, A., Wang, M. C., Veniero, D., & Pons, J. L. (2023). Can we manipulate brain connectivity? A systematic review of cortico-cortical paired associative stimulation effects. Clinical Neurophysiology, 154, 169–193. 10.1016/j.clinph.2023.06.016

47. Hernandez-Pavon, J. C., Veniero, D., Bergmann, T. O., Belardinelli, P., Bortoletto, M., Casarotto, S., Casula, E. P., Farzan, F., Fecchio, M., Julkunen, P., Kallioniemi, E., Lioumis, P., Metsomaa, J., Miniussi, C., Mutanen, T. P., Rocchi, L., Rogasch, N. C., Shafi, M. M., Siebner, H. R., … Ilmoniemi, R. J. (2023). TMS combined with EEG: Recommendations and open issues for data collection and analysis. Brain Stimulation, 16(2), 567–593. 10.1016/J.BRS.2023.02.009

48. Ilmoniemi, R. J., Virtanen, J., Ruohonen, J., Karhu, J., Aronen, H. J., Näätänen, R., & Katila, T. (1997). Neuronal responses to magnetic stimulation reveal cortical reactivity and connectivity. NeuroReport, 8(16), 3537–3540. 10.1097/00001756-199711100-00024

49. Kammer, T., Beck, S., Erb, M., & Grodd, W. (2001). The influence of current direction on phosphene thresholds evoked by transcranial magnetic stimulation. Clinical Neurophysiology, 112(11), 2015–2021. 10.1016/S1388-2457(01)00673-3

50. Kleinert, T., Koenig, T., Nash, K., & Wascher, E. (2024). On the Reliability of the EEG Microstate Approach. Brain Topography, 37(2), 271–286. 10.1007/s10548-023-00982-9

51. Koch, G., Ponzo, V., Di Lorenzo, F., Caltagirone, C., & Veniero, D. (2013). Hebbian and Anti-Hebbian Spike-Timing-Dependent Plasticity of Human Cortico-Cortical Connections. Journal of Neuroscience, 33(23), 9725–9733. 10.1523/JNEUROSCI.4988-12.2013

52. Koenig, T., Kottlow, M., Stein, M., & Melie-García, L. (2011). Ragu: A free tool for the analysis of EEG and MEG event-related scalp field data using global randomization statistics. Computational Intelligence and Neuroscience, 2011. 10.1155/2011/938925

53. Koenig, T., Stein, M., Grieder, M., & Kottlow, M. (2014). A tutorial on data-driven methods for statistically assessing ERP topographies. Brain Topography, 27(1), 72–83. 10.1007/s10548-013-0310-1

54. Lehmann, D., Ozaki, H., & Pal, I. (1987). EEG alpha map series: brain micro-states by space-oriented adaptive segmentation. Electroencephalography and Clinical Neurophysiology, 67(3), 271–288. 10.1016/0013-4694(87)90025-3

55. Luscher, B., Shen, Q., & Sahir, N. (2011). The GABAergic deficit hypothesis of major depressive disorder. In Molecular Psychiatry (Vol. 16, Issue 4, pp. 383–406). 10.1038/mp.2010.120

56. Maccabee, P. J., Nagarajan, S. S., Amassian, V. E., Durand, D. M., Szabo, A. Z., Ahad, A. B., Cracco, R. Q., Lai, K. S., & Eberle, L. P. (1998). Influence of pulse sequence, polarity and amplitude on magnetic stimulation of human and porcine peripheral nerve. Journal of Physiology, 513(2), 571–585. 10.1111/j.1469-7793.1998.571bb.x

57. Mair, P., & Wilcox, R. (2020). Robust statistical methods in R using the WRS2 package. Behavior Research Methods, 52(2), 464–488. 10.3758/s13428-019-01246-w

58. Mäki, H., & Ilmoniemi, R. J. (2010). The relationship between peripheral and early cortical activation induced by transcranial magnetic stimulation. Neuroscience Letters, 478(1), 24–28. 10.1016/j.neulet.2010.04.059

59. Mao, T., Kusefoglu, D., Hooks, B. M., Huber, D., Petreanu, L., & Svoboda, K. (2011). Long-Range Neuronal Circuits Underlying the Interaction between Sensory and Motor Cortex. Neuron, 72(1), 111–123. 10.1016/j.neuron.2011.07.029

60. Matelli, M., Luppino, G., Fogassi, L., & Rizzolatti, G. (1989). Thalamic input to inferior area 6 and area 4 in the macaque monkey. Journal of Comparative Neurology, 280(3), 468–488. 10.1002/cne.902800311

61. Michel, C. M., Brechet, L., Schiller, B., & Koenig, T. (2024). Current State of EEG/ERP Microstate Research. Brain Topography, 169–180. 10.1007/s10548-024-01037-3

62. Michel, C. M., & Koenig, T. (2018). EEG microstates as a tool for studying the temporal dynamics of whole-brain neuronal networks: A review. NeuroImage, 180(May 2017), 577–593. 10.1016/j.neuroimage.2017.11.062

63. Mills, K. R., Boniface, S. J., & Schubert, M. (1992). Magnetic brain stimulation with a double coil: the importance of coil orientation. Electroencephalography and Clinical Neurophysiology/Evoked Potentials Section, 85(1), 17–21. 10.1016/0168-5597(92)90096-T

64. Murray, M. M., Brunet, D., & Michel, C. M. (2008). Topographic ERP analyses: A step-by-step tutorial review. Brain Topography, 20(4), 249–264. 10.1007/s10548-008-0054-5

65. Mutanen, T. P., Kukkonen, M., Nieminen, J. O., Stenroos, M., Sarvas, J., & Ilmoniemi, R. J. (2016). Recovering TMS-evoked EEG responses masked by muscle artifacts. NeuroImage, 139, 157–166. 10.1016/j.neuroimage.2016.05.028

66. Mutanen, T. P., Metsomaa, J., Liljander, S., & Ilmoniemi, R. J. (2018). Automatic and robust noise suppression in EEG and MEG: The SOUND algorithm. NeuroImage, 166(October 2017), 135–151. 10.1016/j.neuroimage.2017.10.021

67. Ni, Z., Charab, S., Gunraj, C., Nelson, A. J., Udupa, K., Yeh, I. J., & Chen, R. (2011). Transcranial magnetic stimulation in different current directions activates separate cortical circuits. Journal of Neurophysiology, 105(2), 749–756. 10.1152/jn.00640.2010

68. Oostenveld, R., Fries, P., Maris, E., & Schoffelen, J. M. (2011). FieldTrip: Open source software for advanced analysis of MEG, EEG, and invasive electrophysiological data. Computational Intelligence and Neuroscience, 2011. 10.1155/2011/156869

69. Petrichella, S., Johnson, N., & He, B. (2017). The influence of corticospinal activity on TMS-evoked activity and connectivity in healthy subjects: A TMS-EEG study. PLoS ONE, 12(4), 1–18. 10.1371/journal.pone.0174879

70. Pieramico, G., Guidotti, R., Nieminen, A. E., D’Andrea, A., Basti, A., Souza, V. H., Nieminen, J. O., Lioumis, P., Ilmoniemi, R. J., Romani, G. L., Pizzella, V., & Marzetti, L. (2023). TMS-Induced Modulation of EEG Functional Connectivity Is Affected by the E-Field Orientation. Brain Sciences, 13(3). 10.3390/brainsci13030418

71. Premoli, I., Castellanos, N., Rivolta, D., Belardinelli, P., Bajo, R., Zipser, C., Espenhahn, S., Heidegger, T., Müller-Dahlhaus, F., & Ziemann, U. (2014). TMS-EEG signatures of GABAergic neurotransmission in the human cortex. Journal of Neuroscience, 34(16), 5603–5612. 10.1523/JNEUROSCI.5089-13.2014

72. Premoli, I., Király, J., Müller-Dahlhaus, F., Zipser, C. M., Rossini, P., Zrenner, C., Ziemann, U., & Belardinelli, P. (2018). Short-interval and long-interval intracortical inhibition of TMS-evoked EEG potentials. Brain Stimulation, 11(4), 818–827. 10.1016/j.brs.2018.03.008

73. Premoli, I., Rivolta, D., Espenhahn, S., Castellanos, N., Belardinelli, P., Ziemann, U., & Müller-Dahlhaus, F. (2014). Characterization of GABAB-receptor mediated neurotransmission in the human cortex by paired-pulse TMS-EEG. NeuroImage, 103, 152–162. 10.1016/j.neuroimage.2014.09.028

74. R Core Team. (2020). R: A Language and Environment for Statistical Computing.

75. Rogasch, N. C., Daskalakis, Z. J., & Fitzgerald, P. B. (2013). Mechanisms underlying long-interval cortical inhibition in the human motor cortex: a TMS-EEG study. J Neurophysiol, 109, 89–98. 10.1152/jn.00762.2012.-Long-interval

76. Rogasch, N. C., & Fitzgerald, P. B. (2013). Assessing cortical network properties using TMS-EEG. Human Brain Mapping, 34(7), 1652–1669. 10.1002/hbm.22016

77. Rossi, S., Antal, A., Bestmann, S., Bikson, M., Brewer, C., Brockmöller, J., Carpenter, L. L., Cincotta, M., Chen, R., Daskalakis, J. D., Di Lazzaro, V., Fox, M. D., George, M. S., Gilbert, D., Kimiskidis, V. K., Koch, G., Ilmoniemi, R. J., Pascal Lefaucheur, J., Leocani, L., … Hallett, M. (2021). Safety and recommendations for TMS use in healthy subjects and patient populations, with updates on training, ethical and regulatory issues: Expert Guidelines. Clinical Neurophysiology, 132(1), 269–306. 10.1016/J.CLINPH.2020.10.003

78. Sakai, K., Ugawa, Y., Terao, Y., Hanajima, R., Furubayashi, T., & Kanazawa, I. (1997). Preferential activation of different I waves by transcranial magnetic stimulation with a figure-of-eight-shaped coil. Experimental Brain Research, 113(1), 24–32. 10.1007/BF02454139

79. Sale, M. V., Lavender, A. P., Opie, G. M., Nordstrom, M. A., & Semmler, J. G. (2016). Increased intracortical inhibition in elderly adults with anterior-posterior current flow: A TMS study. Clinical Neurophysiology, 127(1), 635–640. 10.1016/j.clinph.2015.04.062

80. Siebner, H. R., Funke, K., Aberra, A. S., Antal, A., Bestmann, S., Chen, R., Classen, J., Davare, M., Di Lazzaro, V., Fox, P. T., Hallett, M., Karabanov, A. N., Kesselheim, J., Beck, M. M., Koch, G., Liebetanz, D., Meunier, S., Miniussi, C., Paulus, W., … Ugawa, Y. (2022). Transcranial magnetic stimulation of the brain: What is stimulated? – A consensus and critical position paper. Clinical Neurophysiology, 140, 59–97. 10.1016/J.CLINPH.2022.04.022

81. Sommer, M., Alfaro, A., Rummel, M., Speck, S., Lang, N., Tings, T., & Paulus, W. (2006). Half sine, monophasic and biphasic transcranial magnetic stimulation of the human motor cortex. Clinical Neurophysiology, 117(4), 838– 844. 10.1016/j.clinph.2005.10.029

82. Sommer, M., Ciocca, M., Chieffo, R., Hammond, P., Neef, A., Paulus, W., Rothwell, J. C., & Hannah, R. (2018). TMS of primary motor cortex with a biphasic pulse activates two independent sets of excitable neurones. Brain Stimulation, 11(3), 558–565. 10.1016/j.brs.2018.01.001

83. Sommer, M., Norden, C., Schmack, L., Rothkegel, H., Lang, N., & Paulus, W. (2013). Opposite optimal current flow directions for induction of neuroplasticity and excitation threshold in the human motor cortex. Brain Stimulation, 6(3), 363–370. 10.1016/j.brs.2012.07.003

84. Souza, V. H., Vieira, T. M., Peres, A. S. C., Garcia, M. A. C., Vargas, C. D., & Baffa, O. (2018). Effect of TMS coil orientation on the spatial distribution of motor evoked potentials in an intrinsic hand muscle. Biomedizinische Technik, 63(6), 635–645. 10.1515/bmt-2016-0240

85. Spampinato, D. (2020). Dissecting two distinct interneuronal networks in M1 with transcranial magnetic stimulation. Experimental Brain Research, 238(7–8), 1693–1700. 10.1007/s00221-020-05875-y

86. Stepniewska, I., Preuss, T. M., & Kaas, J. H. (1993). Architectionis, somatotopic organization, and ipsilateral cortical connections of the primary motor area (M1) of owl monkeys. Journal of Comparative Neurology, 330(2), 238–271. 10.1002/cne.903300207

87. Stepniewska, I., Preuss, T. M., & Kaas, J. H. (1994). Thalamic connections of the primary motor cortex (M1) of owl monkeys. Journal of Comparative Neurology, 349(4), 558–582. 10.1002/cne.903490405

88. Sulcova, D., Salatino, A., Ivanoiu, A., & Mouraux, A. (2022). Investigating the Origin of TMS-evoked Brain Potentials Using Topographic Analysis. Brain Topography, 35(5–6), 583–598. 10.1007/s10548-022-00917-w

89. Tarailis, P., Koenig, T., Michel, C. M., & Griškova-Bulanova, I. (2024). The Functional Aspects of Resting EEG Microstates: A Systematic Review. In Brain Topography (Vol. 37, Issue 2, pp. 181–217). Springer. 10.1007/s10548-023-00958-9

90. The Jamovi Project. (2023). *Jamovi (version 2.3) [Computer Software]. Retrieved from* https://www.jamovi.org.

91. Tings, T., Lang, N., Tergau, F., Paulus, W., & Sommer, M. (2005). Orientation-specific fast rTMS maximizes corticospinal inhibition and facilitation. Experimental Brain Research, 164(3), 323–333. 10.1007/s00221-005-2253-6

92. Tremblay, S., Rogasch, N. C., Premoli, I., Blumberger, D. M., Casarotto, S., Chen, R., Di Lazzaro, V., Farzan, F., Ferrarelli, F., Fitzgerald, P. B., Hui, J., Ilmoniemi, R. J., Kimiskidis, V. K., Kugiumtzis, D., Lioumis, P., Pascual-Leone, A., Pellicciari, M. C., Rajji, T., Thut, G., … Daskalakis, Z. J. (2019). Clinical utility and prospective of TMS–EEG. Clinical Neurophysiology, 130(5), 802–844. 10.1016/j.clinph.2019.01.001

93. Vaughan, H. G. (1982). THE NEURAL ORIGINS OF HUMAN EVENT-RELATED POTENTIALS. Annals of the New York Academy of Sciences, 388(1), 125–138. 10.1111/j.1749-6632.1982.tb50788.x

94. Weiler, N., Wood, L., Yu, J., Solla, S. A., & Shepherd, G. M. G. (2008). Top-down laminar organization of the excitatory network in motor cortex. Nature Neuroscience, 11(3), 360–366. 10.1038/nn2049

95. Wilks, S. S. (1961). Multivariate Statistical Outliers. In *The Indian Journal of Statistics*, Series A (Vol. 25, Issue 4). https://about.jstor.org/terms

96. Zazio, A., Lanza, C. M., Stango, A., Guidali, G., Marcantoni, E., Lucarelli, D., Meloni, S., Bolognini, N., Rossi, R., & Bortoletto, M. (2024). Investigating visuo-tactile mirror properties in borderline personality disorder: A TMS-EEG study. Clinical Neurophysiology, 168, 139–152. 10.1016/j.clinph.2024.10.014

97. Zazio, A., Barchiesi, G., Ferrari, C., Marcantoni, E., & Bortoletto, M. (2022). M1-P15 as a cortical marker for transcallosal inhibition: A preregistered TMS-EEG study. Frontiers in Human Neuroscience, 16. 10.3389/fnhum.2022.937515

98. Zazio, A., Miniussi, C., & Bortoletto, M. (2021). Alpha-band cortico-cortical phase synchronization is associated with effective connectivity in the motor network. Clinical Neurophysiology, 132(10), 2473–2480. 10.1016/J.CLINPH.2021.06.025

