## Supplemental files for "Stimulation parameters shape effective connectivity pathways: insights from microstate analysis on TMS-evoked potentials"

### - SUPPLEMENTARY MATERIALS -

#### SUPPLEMENTAL FIGURES

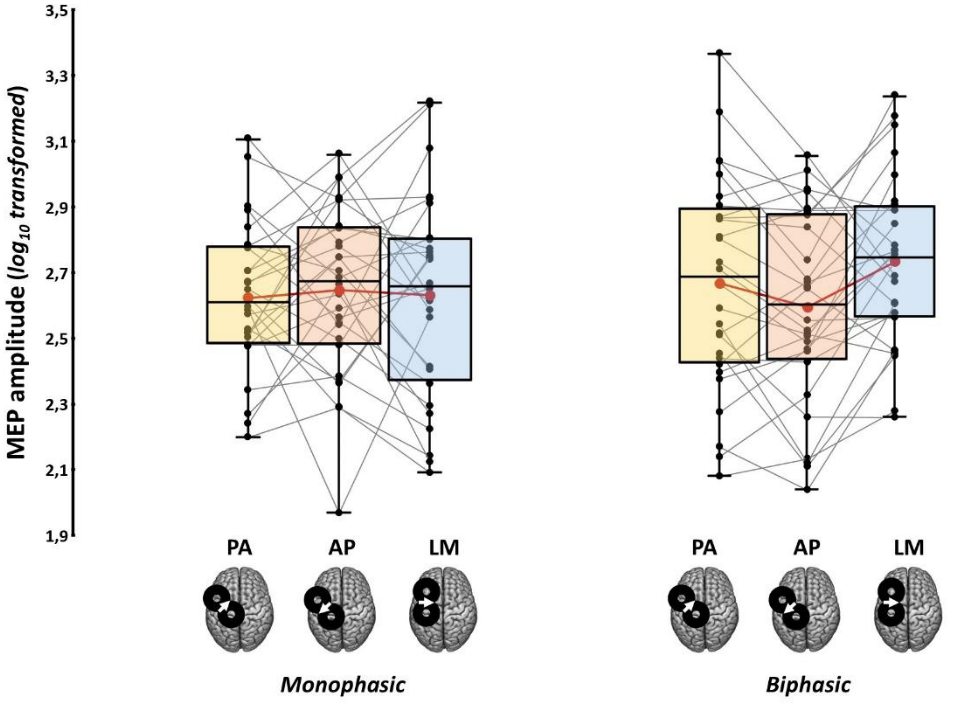


**Supplemental Figure 1.** $Log_{10}$-transformed MEP amplitude recorded in the six experimental conditions in the original study of Guidali et al. (2023). Data were analyzed with a within-subject repeated-measures analysis of variance (rm-ANOVA) with factors 'Pulse waveform' (monophasic, biphasic) and 'Current direction' (PA, AP, LM). No main effect was detected for factors 'Pulse waveform' ($F_{1,27} = 1.39$; $p = .249$; $\eta_p^2 = .049$) and 'Current direction' ($F_{2,54} = 1.46$; $p = .242$; $\eta_p^2 = .051$), and their interaction ($F_{2,54} = 2.76$; $p = .072$; $\eta_p^2 = .093$). In the box-and-whiskers plots, red dots and lines represent the means of the distributions. The center line depicts their median values. Black dots and grey lines show individual scores. The box contains the 25th to 75th percentiles of the dataset. Whiskers extend to the largest observation, which falls within the 1.5 times interquartile range from the first/third quartile. For further information see: Guidali et al., 2023.

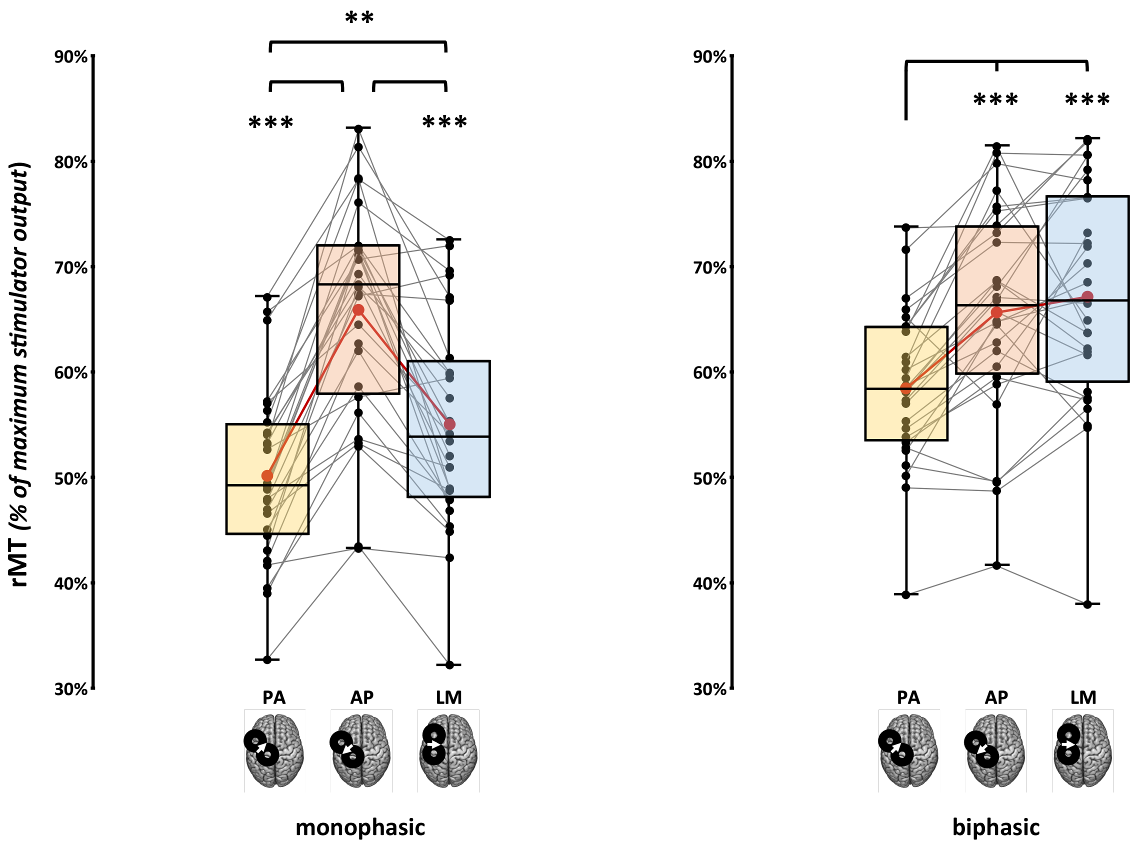


**Supplemental Figure 2.** rMT values recorded in the six experimental conditions in the original study of Guidali et al. (2023). rm-ANOVAs showed significant effects of ‘Current direction’ for both monophasic ($F_{2,54}$ = 60.46; $p < .001$; $\eta_p^2 = .69$) and biphasic conditions ($F_{2,54}$ = 25.44; $p < .001$; $\eta_p^2 = .49$). In the box-and-whiskers plots, red dots and lines represent the means of the distributions. The center line depicts their median values. Black dots and grey lines show individual scores. The box contains the 25th to 75th percentiles of the dataset. Whiskers extend to the largest observation, which falls within the 1.5 times interquartile range from the first/third quartile; significant p-values of corrected Tukey’s post hoc comparisons are reported (** = p <.01; *** = p <.001). For further information see: Guidali et al., 2023.

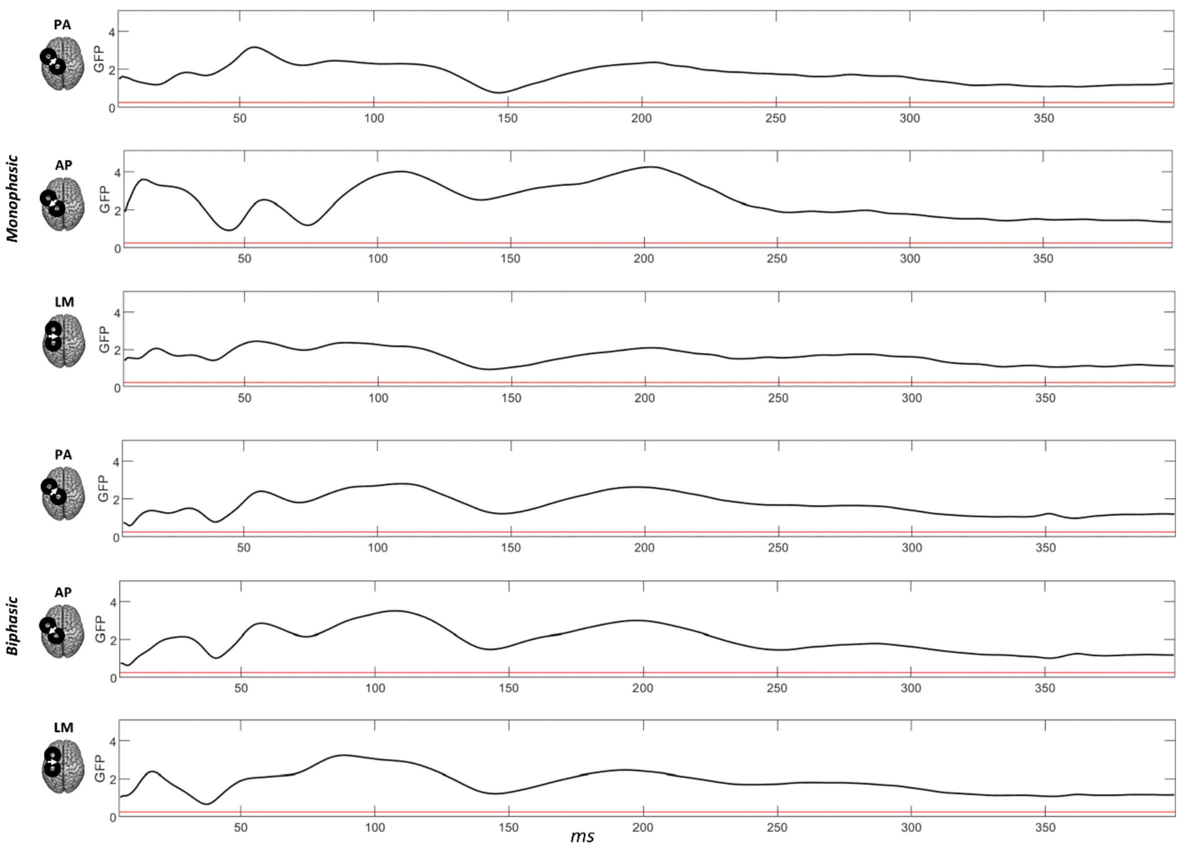


**Supplemental Figure 3.** TCT results. The black line represents the global field power of the grand average signal between 5 and 400 ms post-stimulus for each condition. The red line represents the p-value (=.05).

### SUPPLEMENTAL TABLES

| TEP component | Outliers | F |
|---|---|---|
| **N15** | 3 | 15 |
| **P30** | 3 | 16 |
| **N45** | 3 | 16 |
| **P60** | 5 | 15 |
| **N100** | 3 | 16 |
| **P180** | 3 | 16 |

**Supplemental Table 1.** All the outliers identified for each TEP component. F: number of females.

| *TEP component* | *Factor* | *F* | *p* | $\eta_p^2$ |
|---|---|---|---|---|
| **N15** | **Current direction** | **24.87** | **<.001** | **0.47** |
| | Pulse waveform | 0.23 | .634 | 0.01 |
| | **Current direction X Pulse waveform** | **26.53** | **<.001** | **0.49** |
| **P30** | **Current direction** | **6.42** | **.003** | **0.19** |
| | **Pulse waveform** | **6.12** | **.02** | **0.18** |
| | **Current direction X Pulse waveform** | **11.37** | **<.001** | **0.29** |
| **N45** | **Current direction** | **19.67** | **<.001** | **0.41** |
| | Pulse waveform | 1.39 | .249 | 0.05 |
| | **Current direction X Pulse waveform** | **35.98** | **<.001** | **0.56** |
| **P60** | **Current direction** | **7.14** | **.001** | **0.22** |
| | Pulse waveform | 0.28 | .599 | 0.01 |
| | **Current direction X Pulse waveform** | **5.29** | **0.02** | **0.17** |
| **N100** | **Current direction** | **9.28** | **<.001** | **0.25** |
| | Pulse waveform | 3.31 | .08 | 0.11 |
| | **Current direction X Pulse waveform** | **22.06** | **<.001** | **0.44** |
| **P180** | **Current direction** | **27.61** | **<.001** | **0.5** |
| | Pulse waveform | 0.63 | .434 | 0.02 |
| | **Current direction X Pulse waveform** | **19.7** | **<.001** | **0.41** |

**Supplemental Table 2.** rmANOVAs results conducted for TEP components' amplitudes. Significant main effects and interactions are highlighted in bold.

| *TEP component* | *Factor* | *F* | *p* | $\eta_p^2$ |
|---|---|---|---|---|
| **N15** | **Current direction** | **10.19** | **<.001** | **.27** |
| | **Pulse waveform** | **5.52** | **.026** | **.17** |
| | **Current direction X Pulse waveform** | **10.39** | **<.001** | **.27** |

| *TEP component* | *Pulse waveform* | *Factor* | *F* | *p* |
|---|---|---|---|---|
| **P30** | Monophasic | Current direction | 2.31 | .126 |
| | Biphasic | Current direction | 2.56 | .092 |
| **N45** | Monophasic | Current direction | 0.16 | .849 |
| | Biphasic | Current direction | 1.12 | .338 |
| **P60** | **Monophasic** | **Current direction** | **5.57** | **.013** |
| | Biphasic | Current direction | 1.65 | .21 |
| **N100** | Monophasic | Current direction | 0.29 | .746 |
| | Biphasic | Current direction | 0.86 | .424 |
| **P180** | Monophasic | Current direction | 2.15 | .131 |
| | Biphasic | Current direction | 0.03 | .962 |

**Supplemental Table 3.** rmANOVA (for N15) and robust rmANOVAs results conducted for TEP components' latencies. Significant main effects and interactions are highlighted in bold.

| *Microstate class* | *Pulse waveform* | *Factor* | *F* | *p* |
|---|---|---|---|---|
| **Class 1** | **Monophasic** | **Current direction** | **22.4** | **<.001** |
| | Biphasic | Current direction | 0.65 | .53 |
| **Class 2** | Monophasic | Current direction | 1.38 | .266 |
| | Biphasic | Current direction | 0.03 | .97 |
| **Class 3** | **Monophasic** | **Current direction** | **20.68** | **<.001** |
| | Biphasic | Current direction | 1.55 | .229 |
| **Class 4** | Monophasic | Current direction | 0.36 | .663 |
| | Biphasic | Current direction | 0.89 | .417 |
| **Class 5** | Monophasic | Current direction | 0.26 | .716 |
| | Biphasic | Current direction | 1.07 | .353 |
| **Class 6** | **Monophasic** | **Current direction** | **14.08** | **<.001** |
| | Biphasic | Current direction | 1.33 | .277 |

**Supplemental Table 4.** Robust-rmANOVAs results conducted on microstate AUC. Significant main effects and interactions are highlighted in bold.

| *Microstate class* | *Factor* | *F* | *p* | $\eta_p^2$ |
|---|---|---|---|---|
| **Class 1** | **Current direction** | **15** | **.001** | **0.36** |
| | **Pulse waveform** | **13.5** | **.001** | **0.34** |
| | **Current direction X Pulse waveform** | **16.3** | **<.001** | **0.38** |
| **Class 2** | Current direction | 0.65 | .53 | 0.02 |
| | Pulse waveform | 0.2 | .658 | 0.01 |
| | Current direction X Pulse waveform | 0.53 | .593 | 0.02 |
| **Class 3** | **Current direction** | **19.8** | **<.001** | **0.43** |
| | **Pulse waveform** | **7.67** | **.01** | **0.23** |
| | **Current direction X Pulse waveform** | **18.35** | **<.001** | **0.41** |
| **Class 4** | Current direction | 0.24 | .79 | 0.01 |
| | **Pulse waveform** | **11.74** | **.002** | **0.31** |
| | Current direction X Pulse waveform | 0.01 | .99 | 0.00 |
| **Class 5** | Current direction | 1.04 | .361 | 0.04 |
| | Pulse waveform | 0.06 | .816 | 0,01 |
| | Current direction X Pulse waveform | 1.79 | .176 | 0.07 |
| **Class 6** | **Current direction** | **10.15** | **<.001** | **0.28** |
| | Pulse waveform | 0.03 | .867 | 0.01 |
| | **Current direction X Pulse waveform** | **10.94** | **<.001** | **0.29** |

**Supplemental Table 5.** rmANOVAs results conducted on microstate duration. Significant main effects and interactions are highlighted in bold.
